# Elevated YBX1 mRNA expression is associated with a genomically unstable and clinically aggressive cancer state: a pan-cancer analysis

**DOI:** 10.64898/2026.03.19.712993

**Authors:** Selena Wang, Zahra Shafei Pishabad, Debina Sarkar, Apeksha Bhandarkar, Makhdoom Sarwar, Aaron Jeffs, Glen Reid, Antony Braithwaite, Sunali Mehta

**Affiliations:** Department of Pathology and Molecular Medicine, University of Otago, Dunedin, New Zealand; Maurice Wilkins Centre for Biodiscovery, University of Auckland, Auckland, New Zealand; Department of Obstetrics and Gynaecology, University of Otago, Christchurch, New Zealand; School of Pharmacy and Biomedical Sciences, The University of Waikato Te Whare Wananga o Waikato, Hamilton, New Zealand

**Keywords:** YBX1, YB-1, pan-cancer, genomic instability, chromosomal instability

## Abstract

Y-box binding protein 1 (YB-1; YBX1) is a multifunctional DNA- and RNA-binding protein involved in cell cycle regulation, DNA repair, stress adaptation, and therapy resistance. Elevated YBX1 mRNA expression is associated with aggressive disease across multiple cancers, yet its pan-cancer genomic and clinical correlates remain unclear. Here, we performed a comprehensive pan-cancer analysis across 53 datasets spanning 33 tumour types, integrating RNA expression, somatic mutations, copy number, hypoxia, and clinical outcomes. YBX1 was rarely mutated or amplified, indicating that oncogenic relevance is primarily driven by its expression. Tumours with high YBX1 mRNA exhibited a conserved transcriptional program enriched for cell cycle, DNA repair, and chromatin regulation pathways, and were preferentially mutated in genes involved in maintaining genomic stability, including TP53. These tumours were associated with increased mutation burden, fraction of genome altered, homologous recombination deficiency, and elevated hypoxia. Clinically, high YBX1 mRNA associated with advanced stage, higher grade, shorter progression-free survival, and reduced overall survival. Collectively, high YBX1 mRNA expression defines a conserved, genomically unstable, and clinically aggressive tumour state across multiple cancer types.

## 1. Introduction

The Y-box binding protein 1 (YB-1; encoded by the YBX1 gene) is a cold-shock, multifunctional DNA and RNA binding protein that regulates transcription, mRNA splicing, translation, and cellular stress responses (reviewed in [1], [2], [3], [4]. YB-1 has been implicated in numerous oncogenic processes across multiple cancer types, including regulation of cell cycle progression from the G1/S transition through mitosis and cytokinesis [5], [6], [7], [8], [9], [10], [11], [12], [13], [14]. It also plays a critical role in DNA damage repair, including mismatch repair and double-strand break repair, thereby enabling tolerance to replication stress [15], [16], [17], [18], [19].

In the context of epithelial–mesenchymal transition (EMT), YB-1 acts downstream of Twist to promote the expression of EMT markers such as Snail and vimentin [7], [20], [21], [22]. In addition, YB-1 regulates mRNA splicing of key genes including CD44, which promotes EMT-associated plasticity and invasion, and KLF5, which reinforces EMT-associated transcriptional programs [23], [24]. Effects on EMT are mediated in part through translational activation of HIF1α, a key regulator of hypoxic responses [25], modulation of IL-6 expression [26], and targeting of the intrinsic PD-1/PD-L1 pathway [27].

Furthermore, YB-1 contributes to therapeutic resistance through transcriptional regulation of multi-drug resistance transporters-1 (MDR-1) such as ABCB1 [28], [29], [30], activation of the MDM2/p53 axis [31], [32], suppression of EGR1 expression [33], and activation of AKT signalling [34]. Clinically, elevated YB-1 expression and aberrant subcellular localization are consistently associated with aggressive disease and poor prognosis across a broad spectrum of malignancies [1], [2], [8], [30], underscoring its central role in cancer biology.

A central conceptual question emerging from these observations is how a single gene such as YBX1 can be implicated in such a broad spectrum of oncogenic phenotypes. Rather than independently driving each process, YB-1 may perturb one or more fundamental cellular programs. In a context-dependent manner, these perturbations could result in diverse downstream consequences shaped by tumour lineage, microenvironmental stress, and co-occurring genetic alterations.

A compelling candidate for such a unifying process is genomic instability. Genomic instability can simultaneously promote tumour evolution, phenotypic plasticity, metastatic competence, immune evasion, and therapeutic resistance (reviewed in [35], [36], [37], [38], [39]. Its hallmarks include elevated tumour mutation burden, widespread copy number alterations reflected by an increased fraction of the genome altered. These features arise from defects in DNA damage repair, chromosome segregation, and cell cycle checkpoint control, often facilitated by loss of key tumour suppressors such as *TP53* [35], [40].

Importantly, genomic instability is closely intertwined with tumour hypoxia, a common feature of advanced tumours, with hypoxia stress both driving and resulting from genome destabilisation, thereby promoting the selection of aggressive tumour clones [37]. Although YB-1 has been experimentally linked to cell cycle defects, DNA damage responses, replication stress, and hypoxia-associated signalling [41], it remains unknown whether these processes converge to define a conserved YB-1–associated tumour state across cancers.

Recent studies have begun to examine *YBX1* in a pan-cancer context. The YBX gene family comprises of three closely related Y-box–binding proteins, *YBX1, YBX2*, and *YBX3*, which share a conserved cold shock domain but exhibit distinct expression patterns and biological functions. Pan-cancer analyses of the YBX gene family have demonstrated widespread overexpression of *YBX1* across tumour types, along with associations with oncogenic signalling pathways, immune microenvironment features, and adverse clinical outcomes [42]. Additional large-scale survival analyses have confirmed that high *YBX1* expression is associated with poor overall survival across multiple solid tumours, while other bioinformatic studies have linked *YBX1* expression to immune infiltration and metabolic pathway activity in selected cancers [43], [44]. Collectively, these studies establish *YBX1* as a pan-cancer–relevant gene with prognostic significance.

To date, however, existing analyses have largely focused on differential expression and outcome correlations, leaving fundamental questions unresolved. It remains unclear whether YBX1 is commonly altered at the genomic level or whether its oncogenic relevance is primarily driven by changes in gene dosage and expression. Moreover, despite extensive experimental evidence linking YB-1 to diverse cellular processes, it remains unclear whether elevated YB-1 expression defines a conserved transcriptional program across cancers or instead marks a distinct, clinically aggressive tumour state with shared genomic hallmarks, including genomic instability.

Here, we performed a comprehensive pan-cancer analysis across 53 independent cancer datasets using processed RNA-sequencing gene expression and matched somatic mutation data downloaded from cBioPortal [45], [46], [47]. By leveraging this large and diverse transcriptomic compendium, we show that *YBX1* is infrequently altered at the genomic level across cancers. We identify a conserved *YBX1* mRNA associated gene expression network shared across most cancer types and demonstrate that tumours with high *YBX1* mRNA expression are enriched for mutations in *TP53* and genes involved in genomic and chromosomal stability, exhibit increased tumour mutation burden and fraction of genome altered, and are associated with advanced stage, high grade, elevated hypoxia scores, and poor clinical outcome.

## 2. Results

### 2.1. Somatic alteration and expression patterns of *YBX1* in pan-cancer datasets

Oncogenes are frequently driven by changes in copy number or mutations [48]; therefore, we sought to determine whether *YBX1* follows a similar pattern across cancers. Analysis of multiple somatic pan-cancer datasets, including the TCGA, CPTAC, and BCGSC, encompassing over 12,000 samples across 33 tumour types from 53 studies revealed that YBX1 is largely unaltered, with amplifications occurring in only ∼0.5–8% of cases and mutations in less than ∼4% overall (Figure 1A–C, Supplementary Figure S1).

**Figure 1.**
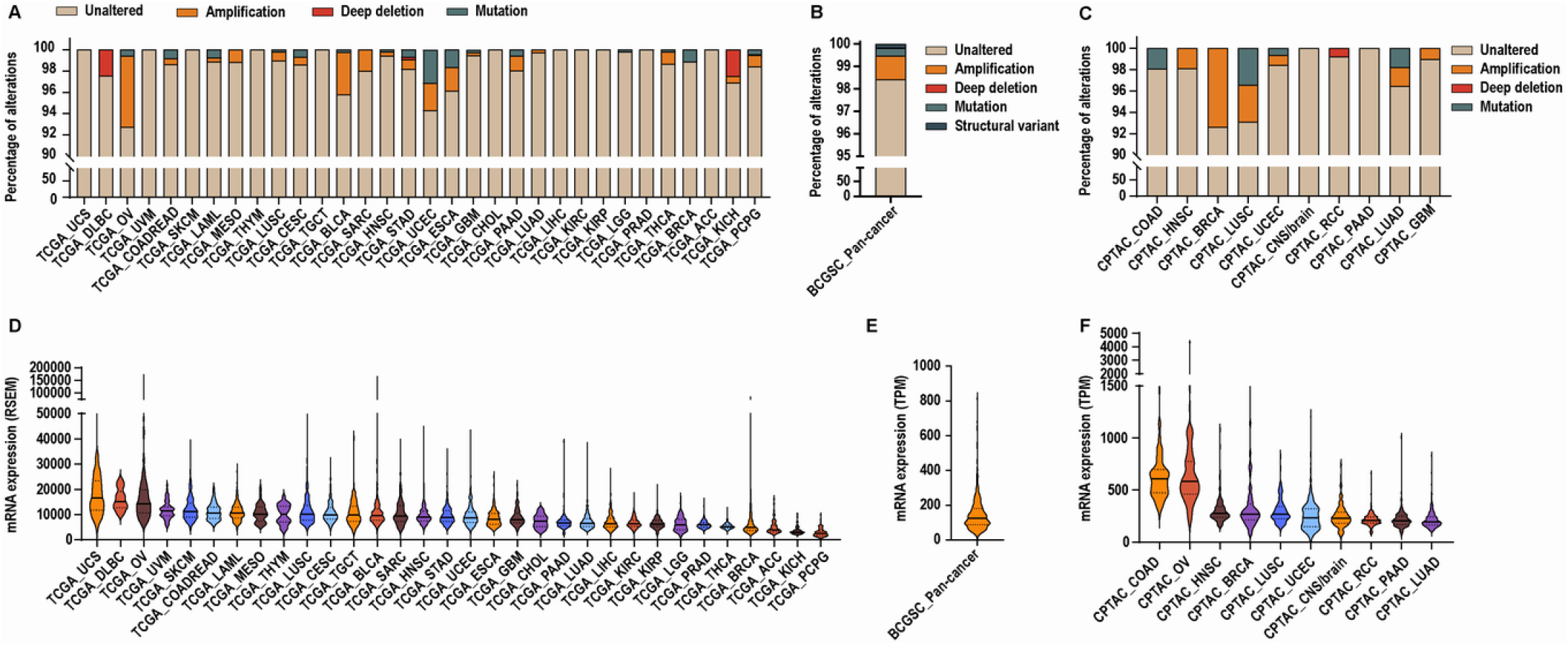
*YBX1* shows broad mRNA expression variability despite minimal genomic alterations across cancers. **A–C**. Bar graphs showing the percentage of alterations in *YBX1* gene from tumour samples within A. Individual cancer types from the TCGA dataset, n = 10,198; **B**. BCGSC pan-cancer study, n = 570; and **C**. individual cancer types from the CPTAC pan-cancer study, n =1,677. **D-F**. Violin plots showing the distribution of *YBX1* mRNA expression across each study. Each violin represents the density of expression values within a tumour group, with the width indicating frequency. The median expression is shown as a dark solid horizontal line, and interquartile range is indicated by dotted lines. **D**. Individual studies from the TCGA dataset, n = 9,367; **E**. BCGSC pan-cancer study, n = 570; and **F**. Individual studies from the CPTAC dataset, n = 1,956.

A few tumour types exhibited slightly higher rates of alteration. In TCGA cohorts, YBX1 was amplified in 6.65% of ovarian cancers (TCGA_OV), 3.93% of bladder cancers (TCGA_BLCA), 2.54% of endometrial cancers (TCGA_UCEC), and 2.2% of oesophageal cancers (TCGA_ESCA). Corresponding mutation frequencies were 3.13% in TCGA_UCEC and 1.65% in TCGA_ESCA cohorts. In CPTAC cohorts, amplification of *YBX1* was observed in 7.37% of breast cancers (CPTAC_BRCA), whereas no amplifications were reported in the corresponding TCGA_BRCA dataset. Additionally, ∼3% of CPTAC_LUSC tumours harboured *YBX1* amplifications and another ∼3% contained *YBX1* mutations, compared with less than 1% of altered cases in the TCGA_LUSC cohort. Despite these dataset- and tumour-specific differences, direct genetic alterations in *YBX1* are rare overall.

In contrast, pan-cancer transcriptomic analyses demonstrated that YBX1 mRNA is broadly expressed across all tumour types but exhibits substantial inter-tumour heterogeneity within cohorts. To quantify this variability, we calculated the expression range within each dataset, defined as the ratio of the maximum to minimum YBX1 mRNA expression observed across tumours in that cohort. Across individual cancer types within the TCGA pan-cancer cohorts, YBX1 mRNA expression spanned expression ranges from 2.5-fold to 135-fold, with a median within-cohort expression range of 10.63-fold. The BCGSC pan-cancer cohort exhibited an expression range of 173-fold, while CPTAC studies showed expression ranges from 5.76-fold to 13,923-fold, with a median expression range of 12.3-fold across studies. These data highlight pronounced heterogeneity of YBX1 mRNA expression within tumour cohorts, supporting the presence of distinct high-YBX1-expressing subsets across diverse cancer lineages (Figure 1D–F, Supplementary Figure S1).

Collectively, these findings indicate that although YBX1 is rarely altered at the genomic level, elevated and heterogeneous expression is a common feature across cancers, suggesting that its oncogenic relevance is primarily mediated through transcriptional or post-transcriptional mechanisms rather than frequent genetic alteration.

### 2.2 Pan-cancer correlation analysis reveals conserved transcriptional programs associated with *YBX1* mRNA expression

To determine whether elevated *YBX1* mRNA expression is associated with specific cancer-related biological programs, we performed Spearman’s correlation analyses between *YBX1* mRNA levels and the mRNA expression of 10,830 genes across 53 independent cancer datasets. Genes were selected based on consistent positive association with *YBX1* expression, defined as a Spearman correlation coefficient (ρ) ≥ 0.3 with q ≤ 0.05 in at least ∼80% (42/53) of the datasets analyzed. This analysis identified 23 genes meeting these criteria (Figure 2A).

**Figure 2.**
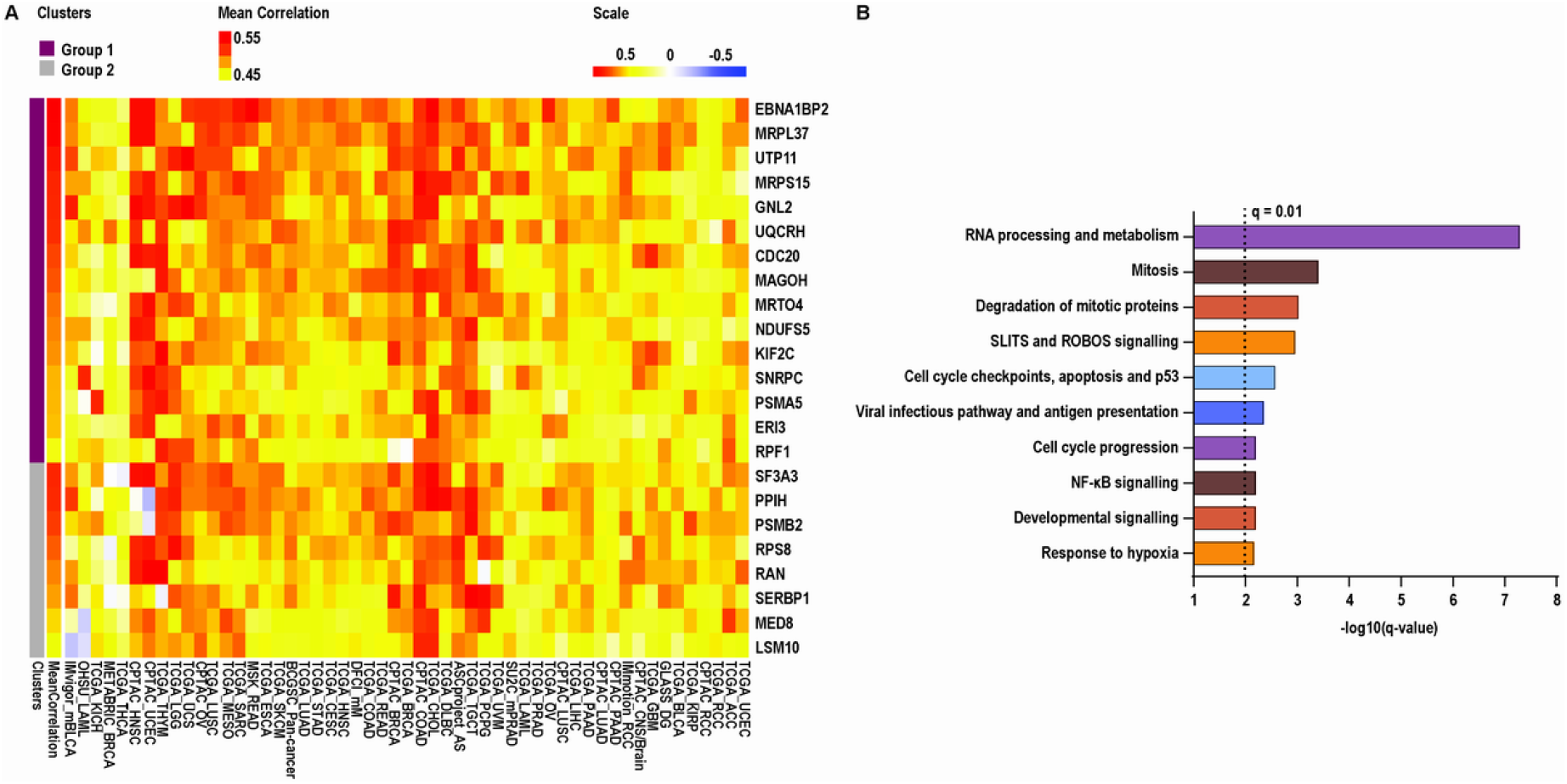
Genes positively correlated with *YBX1* mRNA expression and the top enriched pathways. **A**. Heat map showing genes positively correlated with *YBX1* mRNA in at least 80% of the 53 datasets analysed (42/53). Significance was defined by Spearman’s correlation coefficient ≥ 0.3 and a *q*-value < 0.05. Group 1: Genes clustered in purple (upper quadrant) show exclusively positive correlations with *YBX1*. Group 2: Genes clustered in grey (lower quadrant) display negative correlations with *YBX1* in at least one dataset. Within each cluster, genes are ordered from top to bottom by descending mean correlation across all datasets. **B**. Significantly enriched pathways (q-value ≤ 0.01) identified using EnrichR Reactome Pathway 2024 database for the genes positively correlated with *YBX1* mRNA across datasets.

Notably, 15 of these genes showed positive correlation across all 53 datasets, with significant positive correlations observed in at least 45 datasets, indicating a highly robust association with *YBX1* mRNA expression across diverse tumour contexts. The remaining eight genes were positively correlated in at least 51 of 53 datasets; among these, five genes (*PPIH, PSMB2, RPS8, RAN, MED8*) were negatively correlated in a single dataset, while three genes (*SF3A3, SERBP1*, and *LSM10*) were negatively correlated in two datasets. For visualization purposes, correlation coefficients across all datasets including those not meeting significance thresholds, were displayed in the heatmap (Figure 2A).

In contrast, when the same consistency criteria were applied to negatively correlated genes, none were consistently shared across datasets (ρ ≤ −0.3, q ≤ 0.05 in ∼80% of datasets), underscoring the specificity of positive *YBX1*-associated transcriptional programs.

To assess the biological relevance of these associations, we next performed pathway enrichment analysis on genes whose mRNA expression correlated with *YBX1* mRNA levels. This revealed strong overrepresentation of pathways involved in RNA processing and metabolism, cell cycle regulation at G1/S and G2/M, DNA replication and checkpoint control, hypoxia, and multiple oncogenic signalling pathways (Figure 2B, Supplementary Table S1). Collectively, these findings indicate that high *YBX1* expression is tightly coupled to conserved proliferative and transcriptionally active tumour programs across cancer types.

### 2.3 Pan-Cancer Genomic and Pathway Correlates of *YBX1* Expression

To investigate whether YBX1 expression is associated with specific mutational patterns, we focused our analysis on the TCGA datasets, as they provide the most comprehensive, high-quality, and uniformly processed mutation and expression data across diverse cancer types. Mutation rates for each gene were compared between the high- and low-expression groups, and a delta value (Δ = high − low) was computed to quantify enrichment of mutations in high-*YBX1* tumours. Genes were then classified as Positive (Δ > 0), Negative (Δ < 0), or Unchanged (Δ = 0), and the number of tumour types with positive (pos) versus negative (neg) enrichment was summarized as a pos/neg ratio. Genes with a pos/neg ratio > 2 were considered consistently enriched in high-*YBX1* tumors, whereas those with a ratio ≤ 0.8 were depleted (Figure 3A-B).

**Figure 3.**
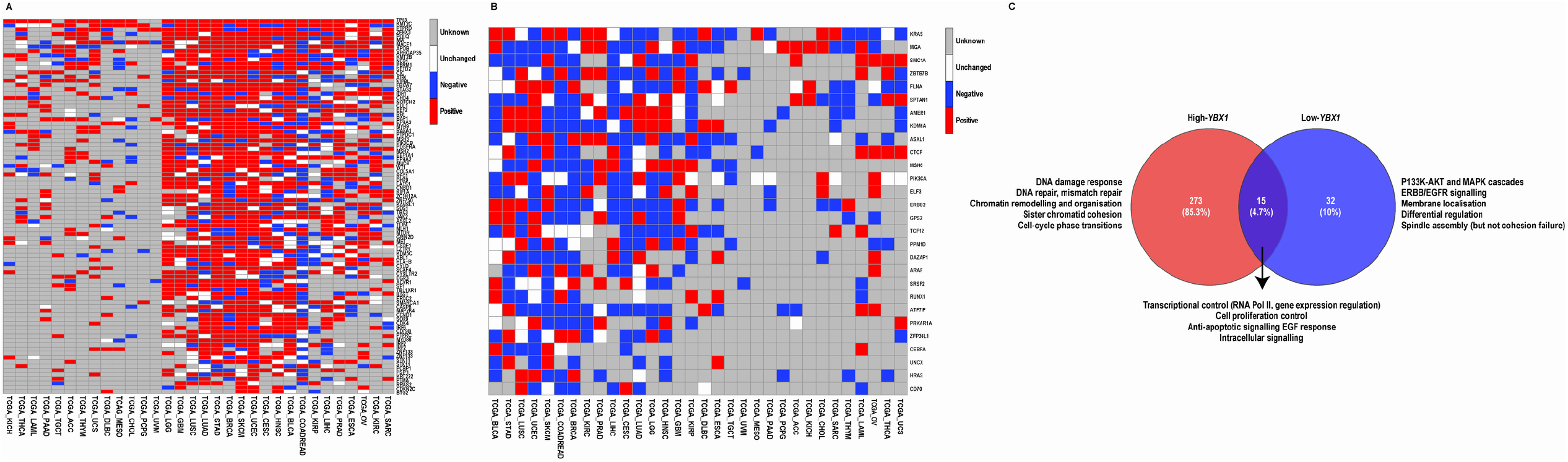
High- and low-*YBX1* tumours display distinct mutation frequencies in known driver genes. **A-B**. Heat maps showing known cancer driver genes that are frequently mutated in **A**. High-*YBX1*, **B**. Low-*YBX1*, tumours across all 32 tumour types in the TCGA dataset. Tumours were stratified into high and low *YBX1* groups based on the 75th and 25th percentiles of *YBX1* mRNA expression within each tumour type. For each gene, mutation enrichment was classified as Positive (red, Δ > 0) if the mutation rate was higher, Negative (blue, Δ < 0) if lower, and Unchanged (white, Δ = 0) if no difference was observed in the high-YBX1 group. Missing data are indicated in grey. Genes were ordered by the number of tumour types with positive enrichment, and hierarchical clustering was applied to both genes and tumour types to reveal patterns of mutation enrichment across datasets. **C**. Venn diagram showing independent and overlapping pathways enriched in the EnrichR GO Biological Processes 2025 database (q-value ≤ 0.01) for genes frequently mutated in high-YBX1, low-YBX1, and shared tumour groups. The major functional classes of enriched GO Biological Processes are shown alongside each group, with full pathway details provided in Supplementary Table S2.

Using this approach, *TP53*, the key tumour suppressor and “guardian of the genome,” was consistently mutated at a higher frequency in high-*YBX1* tumours in 29 of the 32 TCGA datasets analysed (Figure 3A). Beyond *TP53*, multiple genes with well-established roles in maintaining genome integrity were preferentially mutated in high-*YBX1* tumours (Figure 3A). *TP53* was mutated in all datasets where data were available, while the remaining genes were mutated across most datasets. These genes function across multiple genome-stabilizing processes, including DNA repair (*POLQ, SETD2, ATRX, MSH2, MSH3, MLH1, ERCC2, RFC1, DHX9, PSIP1*), chromatin regulation (*KMT2C, KMT2B, NSD1, PBRM1, CHD4, KANSL1, KDM5C, ASXL2, SMARCA1, SCAF4*), chromosome segregation (*STAG2, NIPBL, RB1, LATS1, FBXW7, CUL1*), and cell-cycle checkpoint control (*CDK4, CCND1, CDKN2C*). Collectively, mutations in these pathways are known to promote genomic instability through defects in DNA damage repair, chromatin organization, and mitotic fidelity.

In contrast, low-*YBX1* tumours (Figure 3B) were enriched for mutations in receptor-proximal signalling and line- age-associated genes, including *KRAS, ERBB2, PIK3CA, ARAF, HRAS, ELF3*, and *CEBPA*. Compared with high-YBX1 tumours, this pattern was less uniform across datasets, but is consistent with a more signal-driven, differentiation-associated tumour state (Figure 3B).

To extend these observations beyond individual genes, we performed Gene Ontology based pathway enrichment analysis using cancer driver genes differentially mutated between high and low-*YBX1* tumours. While both groups shared enrichment for core oncogenic programs related to transcriptional regulation, cell proliferation, and growth factor responsiveness, high-*YBX1* tumours showed extensive and unique enrichment for pathways associated with DNA damage response, DNA repair, mismatch repair, chromatin remodelling, chromosome organization, sister chromatid cohesion, and cell-cycle phase transitions (Figure 3C, Supplementary Table S2). These pathway-level signatures are characteristic of replication stress and chromosomal instability, providing functional support for the elevated frequency of genome maintenance gene mutations observed in high-*YBX1* tumours. Conversely, pathways uniquely enriched in low-*YBX1* tumours were largely restricted to PI3K–AKT, MAPK, and ERBB/EGFR signalling cascades, as well as differentiation-related processes, reinforcing the distinction between a genomically unstable, transcriptionally plastic state in high-*YBX1* tumours and a more signal-driven, lineage-restricted state in low-*YBX1* tumours (Figure 3C).

### 2.4 High-*YBX1* Tumours Are Associated with Increased Genomic Instability

To determine whether elevated *YBX1* mRNA expression is associated with increased genomic instability, we next examined whether high-*YBX1* tumours exhibit increased mutational burden, copy number alterations, or defects in homologous recombination (HR), a high-fidelity DNA repair pathway. Tumours were stratified based on *YBX1* mRNA expression, with high-YBX1 tumours defined as those at or above the 75th percentile and low-*YBX1* tumours defined as those at or below the 25th percentile.

We first assessed total number of mutations across the pan-cancer TCGA cohort, the pan-cancer BCGSC dataset, and CPTAC datasets spanning 10 tumour types. Across all datasets, tumours with high *YBX1* mRNA expression exhibited a significantly higher number of somatic mutations compared to low-*YBX1* tumours (Figure 4A).

**Figure 4.**
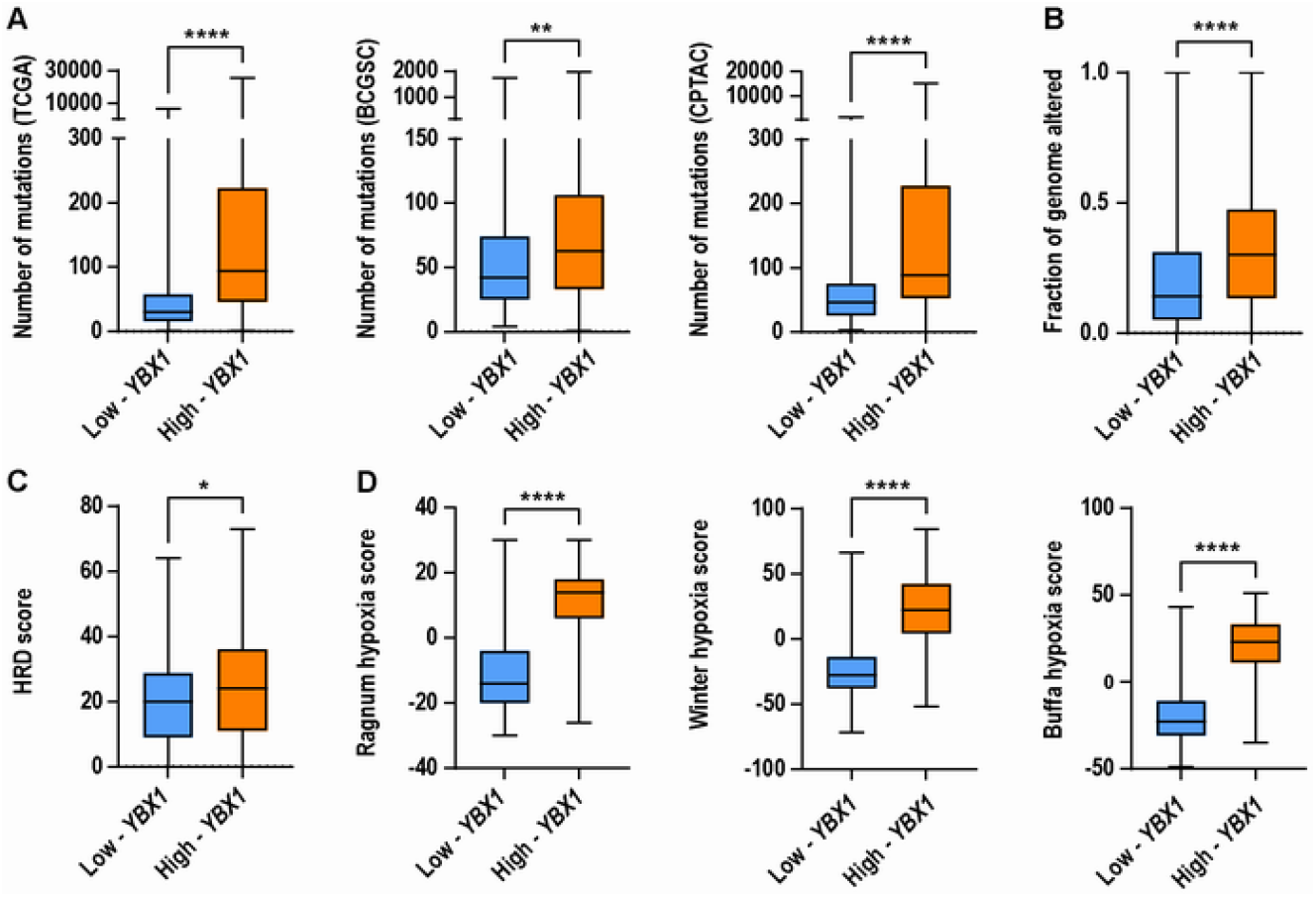
High *YBX1* tumours display increased genomic instability and higher hypoxia score. Box and whiskers plots comparing the distribution of: **A**. Total mutation counts in high- and low-YBX1 tumours across the TCGA (high-*YBX1*, n = 2,360; low-*YBX1*, n = 2,393; ****p < 0.0001), BCGSC (high-*YBX1*, n = 142; low-*YBX1*, n = 143;, **p = 0.0015), and CPTAC (high-*YBX1*, n = 483; low-*YBX1*, n = 278;, ****p < 0.0001) datasets. Significance: Mann-Whitney U test, a p < 0.05 is considered statistically significant. **B**. FGA in high-*YBX1* (n = 2,360) and low-*YBX1* (n = 2,393) tumours in the TCGA dataset (****p < 0.0001). Significance: Mann-Whitney U test, a p < 0.05 is considered statistically significant. **C**. HRD score in high-*YBX1* (n = 144) and low-*YBX1*(n = 143) tumours in the BCGSC dataset (*p = 0.0145). Significance: Mann-Whitney U test, a p < 0.05 is considered statistically significant. D. Hypoxia scores in high-*YBX1* (n = 1722) and low-*YBX1* (n = 2171) tumours in the TCGA dataset (Ragnum: ****p < 0.0001; Buffa: ****p < 0.0001; and Winter: ****p < 0.0001). Significance: Wilcoxon test, a p < 0.05 is considered statistically significant. **A – D**. The central line represents the median, the box indicates the interquartile range (25th–75th percentile), and the whiskers extend the most extreme data point within this range.

Given that genomic instability often manifests not only as point mutations but also as large-scale chromosomal alterations, we next evaluated copy number alterations using the fraction of genome altered (FGA), available for the pan-cancer TCGA cohort. Consistent with increased chromosomal instability, high-*YBX1* tumours displayed significantly higher FGA scores compared to low-YBX1 tumours (Figure 4B).

We further investigated whether high-*YBX1* tumours were associated with impaired homologous recombination repair, available for the pan-cancer BCGSC dataset. This analysis revealed that metastatic tumours with high-*YBX1* expression were significantly more likely to exhibit homologous recombination deficiency compared to their low-*YBX1* counterparts (Figure 4C).

As hypoxia is a well-established feature of genomically unstable tumours and a known contributor to replication stress and DNA repair defects, we next examined the relationship between *YBX1* mRNA expression and tumour hypoxia. Using hypoxia scores available for the pan-cancer TCGA dataset, we found that high-*YBX1* tumours were associated with significantly elevated Ragnum, Winter, and Buffa hypoxia scores relative to low-*YBX1* tumours (Figure 4D).

Taken together, these findings demonstrate that high-*YBX1* mRNA expression is consistently associated with increased mutational burden, chromosomal instability, homologous recombination deficiency and tumour hypoxia, supporting a model in which high-*YBX1* tumours represent a genomically unstable and biologically aggressive tumour state across cancer types.

### 2.5 High *YBX1* mRNA Expression Is Associated with Advanced Tumour Stage, Higher Grade, and Poor Clinical Outcomes

To determine whether the genomic instability associated with high-*YBX1* mRNA expression translates into more aggressive clinical behavior, we next examined the relationship between *YBX1* mRNA expression and key clinicopathological parameters used in patient management. Tumours were stratified into high- and low-*YBX1* groups based on YBX1 mRNA expression (High ≥75th and Low ≤25th percentiles, respectively), and analyses were performed using clinical annotations available from the pan-cancer TCGA dataset.

Consistent with the molecular features observed in high-*YBX1* tumours, we found that these tumours were significantly enriched for advanced disease stage. Specifically, high-*YBX1* tumours exhibited a significantly higher proportion of AJCC T3 and T4 stage disease compared to low-*YBX1* tumours (Figure 5A). In parallel, histopathological assessment revealed that histologic grade was also significantly higher in high-*YBX1* tumours relative to low-*YBX1* tumours, further supporting a more aggressive tumour phenotype (Figure 5B).

**Figure 5.**
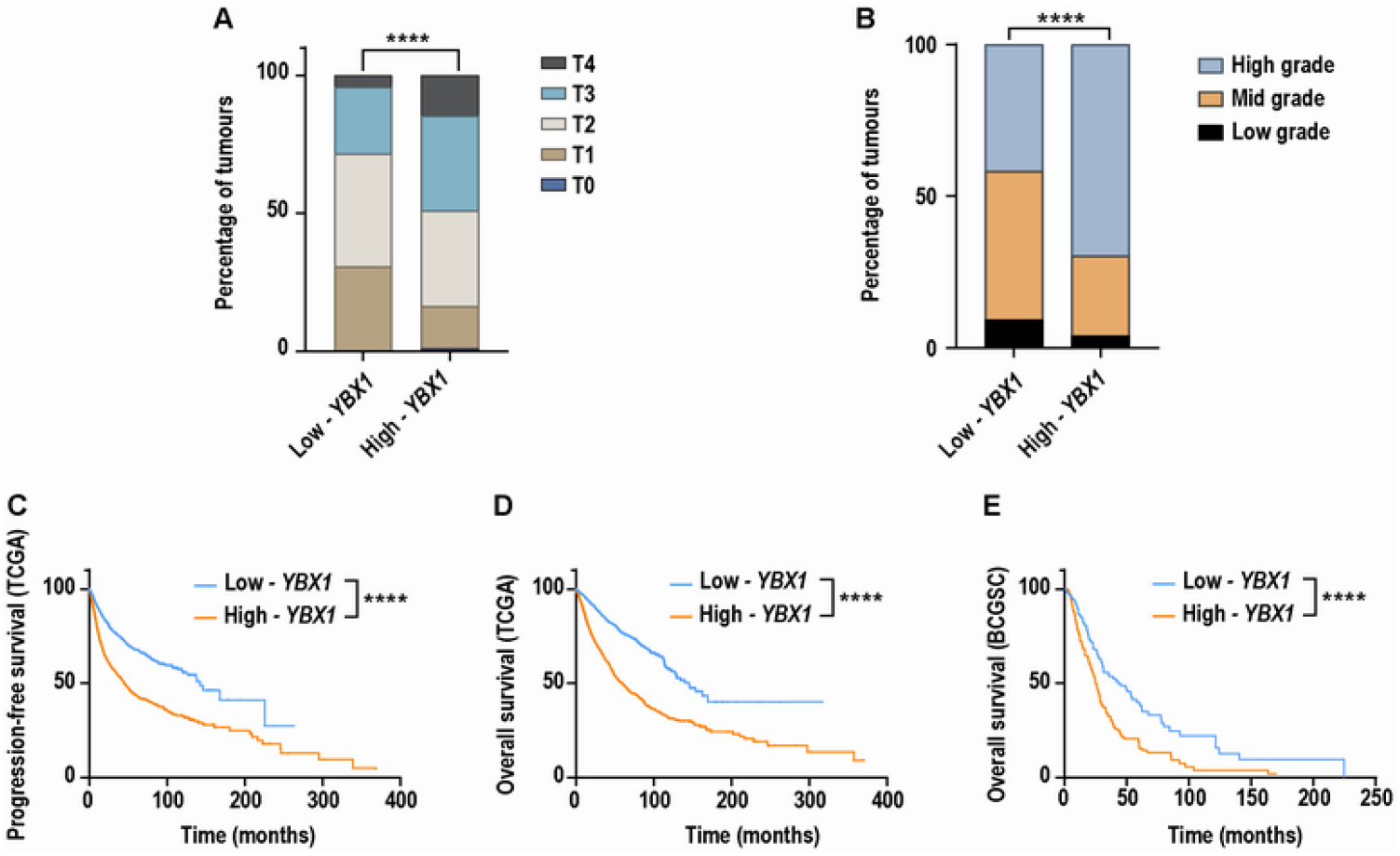
Patients with high *YBX1* mRNA expression exhibit more advanced tumour stage and grade, with reduced survival. Bar graph comparing the percentage distribution of **A**. AJCC tumour stage in patients with high-*YBX1* (n = 1,569) and low-*YBX1* (n = 1,987) tumours across the TCGA dataset (****p < 0.0001). Significance: Chi-squared test, a p < 0.05 is considered statistically significant. T0/TIS: primary tumour not found or cannot be measured; T1-4: increasing in tumour size and invasion into nearby tissues. B. Neoplasm histologic grade in patients with high-*YBX1* (n = 1,027) and low-*YBX1* (n = 684) tumours across the TCGA dataset (****p < 0.0001). Significance: Chi-squared test, p < 0.05 is considered statistically significant. Low grade: well-differentiated tumour cells; mid grade: moderately-differentiated tumour cells; high grade: poorly-differentiated to undifferentiated tumour cells. C-E. Kaplan-Meier plots comparing, **C**. Progression-free survival, TCGA dataset: high-*YBX1* (n = 2,399) and low-*YBX1* (n = 2,511). **D**. Overall survival of patients, TCGA dataset: high-*YBX1* (n = 2,481) and low-*YBX1* (n = 2,514). **E**. Overall survival of patients, BCGSC dataset: high-*YBX1* (n = 143) and low-*YBX1* (n = 144). C-E: Significance: log-rank test, p < 0.05 is considered statistically significant, ****p < 0.0001.

We next evaluated whether elevated *YBX1* mRNA expression was associated with earlier disease recurrence. Progression-free survival was available for the pan-cancer TCGA cohort. This analysis demonstrated that patients with high-*YBX1* tumours experienced a significantly shorter progression-free survival compared to those with low-*YBX1* tumours (Figure 5C).

Finally, to assess whether high-*YBX1* expression also predicts poor outcomes in advanced disease, we analysed overall survival in the pan-cancer TCGA cohort and the metastatic pan-cancer BCGSC dataset. Consistent with the aggressive biology associated with high-*YBX1* tumours, patients with tumours expressing high *YBX1* mRNA exhibited significantly worse overall survival compared to patients whose tumours expressed low levels of *YBX1* mRNA (Figure 5D-E).

Taken together, these findings demonstrate that high *YBX1* mRNA expression is associated with advanced tumour stage, higher histologic grade, earlier disease progression, and reduced survival, linking the genomic instability of *YBX1*-high tumours to clinically aggressive behaviour across cancer types.

## 3. Discussion

This study provides the first comprehensive pan-cancer integration of transcriptomic, genomic, and clinical data to defines a high-YBX1 tumour state conserved across 53 independent datasets spanning 33 tumour types. We identify a reproducible gene expression program associated with elevated *YBX1* mRNA expression, encompassing genomic instability, mitotic dysregulation, DNA repair defects, RNA processing and metabolic alterations, and aggressive clinical features. The robustness of this program across diverse datasets is notable given the known heterogeneity of sequencing platforms [49], [50], the presence of batch effects [51], and variability in clinical annotation across institutions, countries, and time periods [52].

Previous studies have linked elevated YBX1 mRNA or protein expression to poor prognosis [2], [44], [53], [8], therapy resistance [28], [29], [30], [31], [33], [34], and EMT [7], [20], [21], [22] in individual tumour types. Our analysis unifies these observations by demonstrating that high-*YBX1* tumours share conserved molecular and clinical characteristics across multiple cancers, supporting the existence of a pan-cancer YBX1-driven tumour state.

At the genomic level, high-*YBX1* tumours are enriched for *TP53* mutations, suggesting cooperative effects in driving genomic instability and aggressive behaviour. Mechanistically, YB-1 overexpression has been shown to antagonize p53-mediated tumour suppression through MDM2 activation and repression of p53 target genes [31], [32]. Beyond *TP53*, we observe preferential mutation of genes involved in DNA repair, chromatin organization, chromosome segregation, and cell-cycle progression, processes central to genome maintenance. Consistent with these findings, experimental studies have established roles for YB-1 in cell-cycle checkpoint control [5], [8], [9], [14], mitotic fidelity [10], [11], [13], and DNA damage repair [15], [16], [17], [18], [19]. Together, mitotic errors and loss of p53-mediated surveillance are likely to exacerbate chromosomal instability, promoting aneuploidy, structural chromosomal aberrations, and aggressive tumour phenotypes.

Pathway-level analyses further reveal enrichment for replication stress, mitotic checkpoint dysregulation, RNA processing, and ribosome biogenesis programs in high-*YBX1* tumours. These findings align with YB-1’s established functions in RNA splicing, mRNA translation, and transcriptional regulation of oncogenic pathways, including MDM2- and HIF1α-driven stress responses [5], [8], [9], [10], [11], [12],[13], [14], [25], [31], [32], [33], [34]. Collectively, these data support a model in which YB-1 overexpression drives coordinated defects in genome maintenance and transcriptional plasticity, facilitating adaptation to hypoxia and replication stress and underpinning the aggressive phenotype of high-*YBX1* tumours.

Importantly, extensive mechanistic literature already establishes YB-1’s roles in multiple aspects of tumour biology. Building on this foundation, future studies using inducible, CRISPR-based, organoid, and *in vivo* models will be essential to define context-specific dependencies, dissect interactions between YB-1 and *TP53*-mediated genome surveillance, and identify therapeutic vulnerabilities. In particular, targeting replication stress responses, mitotic check- point machinery, RNA processing pathways, or DNA repair dependencies may prove effective in high-*YBX1, TP53*-deficient tumours.

Consistent with these molecular features, high-YBX1 tumours are associated with advanced stage, higher histologic grade, elevated hypoxia scores, shorter progression-free survival, and poorer overall survival in metastatic disease [2], [44], [53], [8]. In contrast, low-*YBX1* tumours preferentially activate receptor-proximal signalling and differentiation-associated pathways, reflecting a more lineage-restricted, signal-driven state. These integrated molecular and clinical observations provide mechanistic context for prior reports implicating YB-1 in EMT, stemness, and chemoresistance across multiple cancer types [7], [20], [21], [22], [23], [24], [28], [29], [30].

Several limitations should be considered. Our analyses rely on bulk transcriptomic and genomic data, precluding resolution of cell type–specific effects and causal inference. Data were generated across multiple institutions, time periods, and platforms, introducing potential batch effects, and clinical annotations were not uniformly available. In addition, our stratification is based on *YBX1* mRNA expression and percentile thresholds, which may vary between tumour types. Moreover, *YBX1* mRNA levels are likely shaped by multiple regulatory inputs, including oncogenic signalling and cellular stress responses, as well as post-transcriptional control mechanisms, and therefore may not fully reflect functional YB-1 protein activity. Nevertheless, the consistent emergence of a *high*-YBX1 molecular program across diverse datasets supports its biological and clinical relevance.

## 4. Materials and Methods

### 4.1. Data retrieval from the cBioPortal Database

A total of 53 datasets from cBioPortal (accessed in January 2026) [45], [46], [47] were downloaded and integrated to analyse the genomic, transcriptomic, and clinical characteristics associated with YBX1 mRNA. This included two pan-cancer datasets, the British Columbia Genome Sciences Centre Pan-cancer Analysis of Advanced and Metastatic Tumors (BCGSC, Nature Cancer 2020), and the Cancer Genome Atlas PanCancer Atlas studies (TCGA, PanCancer Atlas); 16 studies of various cancer types harmonised by the Clinical Proteomic Tumor Analysis Consortium (CPTAC), including Colon Adenocarcinoma (CPTAC GDC, 2025), Colon Cancer (CPTAC-2 Prospective, Cell 2019), CNS/Brain Cancer (CPTAC GDC, 2025), Glioblastoma (CPTAC, Cell 2021), Head and Neck Carcinoma, Other (CPTAC GDC, 2025), Renal Cell Carcinoma (CPTAC GDC, 2025), Lung Adenocarcinoma (CPTAC, Cell 2020; CPTAC GDC, 2025), Lung Squamous Cell Carcinoma (CPTAC, Cell 2021; CPTAC GDC, 2025), Ovarian Cancer (CPTAC GDC, 2025), Pancreatic Cancer (CPTAC GDC, 2025), Endometrial Carcinoma (CPTAC, Cell 2020), Uterine Endometrioid Carcinoma (CPTAC GDC, 2025), Breast Cancer (CPTAC GDC, 2025), Proteogenomic landscape of breast cancer (CPTAC, Cell 2020); and 9 miscellaneous studies of various cancer types including Metastatic Bladder Urothelial Carcinoma (IMvigor210 Phase II Trial, ESMO Open. 2024), Renal Cell Carcinoma (IMmotion150 Clinical Trial, Nat Med. 2018), Metastatic Melanoma (DFCI, Nat Med. 2019), Rectal Cancer (MSK, Nature Medicine 2022), Breast Cancer (METABRIC, Nature 2012 & Nat Commun 2016), Diffuse Glioma (GLASS Consortium), Acute Myeloid Leukemia (OHSU, Nature 2018), Metastatic Prostate Adenocarcinoma (SU2C/PCF Dream Team, PNAS 2019), The Angiosarcoma Project - Count Me In (Provisional, April 2025).

The BCGSC Nature Cancer 2020 dataset (referred to as BCGSC) was retrieved at the pan-cancer level.

The TCGA PanCancer Atlas dataset (referred to as TCGA) was retrieved both at the pan-cancer level and across 33 individual cancer types. These are: uterine carcinosarcoma (TCGA_UCS), diffuse large B-cell lymphoma (TCGA_DLBC), ovarian serous cystadenocarcinoma (TCGA_OV), uveal melanoma (TCGA_UVM), skin cutaneous melanoma (TCGA_SKCM), acute myeloid leukemia (TCGA_LAML), colorectal adenocarcinoma (TCGA_COADREAD; comprising colon adenocarcinoma (TCGA_COAD) and rectal adenocarcinoma (TCGA_READ)), mesothelioma (TCGA_MESO), thymoma (TCGA_THYM), lung squamous cell carcinoma (TCGA_LUSC), cervical squamous cell carcinoma (TCGA_CESC), testicular germ cell tumors (TCGA_TGCT), bladder urothelial carcinoma (TCGA_BLCA), sarcoma (TCGA_SARC), head and neck squamous cell carcinoma (TCGA_HNSC), stomach adeno- carcinoma (TCGA_STAD), uterine corpus endometrial carcinoma (TCGA_UCEC), esophageal adenocarcinoma (TCGA_ESCA), glioblastoma multiforme (TCGA_GBM), cholangiocarcinoma (TCGA_CHOL), pancreatic adenocarcinoma (TCGA_PAAD), lung adenocarcinoma (TCGA_LUAD), liver hepatocellular carcinoma (TCGA_LIHC), kidney renal clear cell carcinoma (TCGA_KIRC), kidney renal papillary cell carcinoma (TCGA_KIRP), brain lower grade glioma (TCGA_LGG), prostate adenocarcinoma (TCGA_PRAD), thyroid carcinoma (TCGA_THCA), breast invasive carcinoma (TCGA_BRCA), adrenocortical carcinoma (TCGA_ACC), kidney chromophobe (TCGA_KICH), pheochromocytoma and paraganglioma (TCGA_PCPG).

Data from 16 CPTAC-harmonised studies (referred to as CPTAC) across 11 individual cancer types were retrieved, including colon cancer (CPTAC_COAD), ovarian cancer (CPTAC_OV), head and neck carcinoma (CPTAC_HNSC), breast cancer (CPTAC_BRCA), lung squamous cell carcinoma (CPTAC_LUSC), endometrial carcinoma (CPTAC_UCEC), CNS/brain Cancer (CPTAC_CNS/brain), renal cell carcinoma (CPTAC_RCC), pancreatic cancer (CPTAC_PAAD), lung adenocarcinoma (CPTAC_LUAD), glioblastoma (CPTAC_GBM).

Data from 9 miscellaneous studies across 9 individual cancer types were retrieved, including Metastatic Bladder Urothelial Carcinoma (IMvigor_mBLCA), Renal Cell Carcinoma (IMmotion_RCC), Metastatic Melanoma (DFCI_mM), Rectal Adenocarcinoma (MSK_READ), Breast Cancer (METABRIC_BRCA), diffuse glioma (GLASS_DG), Acute Mye- loid Leukaemia (OHSU_LAML), metastatic prostate adenocarcinoma (SU2C_mPRAD), Angiosarcoma (ASCproject_AS).

### 4.2 Genomic and Transcriptomic profiles of YBX1

Genomic and transcriptomic profiles of *YBX1*, along with their corresponding tumour origins, were retrieved by querying the *YBX1* gene in cBioPortal across all datasets.

Genomic alterations assessed include mutations (missense, nonsense, truncation, frameshift), copy number variations (amplifications and deletions), and structural variations. Mutation and copy number variation data were available for all BCGSC, TCGA, and CPTAC datasets, whereas structural variation data were only available in the BCGSC and TCGA datasets. For genomic alterations, the BCGSC dataset was assessed at the pan-cancer level, the TCGA dataset was assessed across 33 individual cancer types, the CPTAC dataset was assessed across 10 out of 11 individual cancer types due to data availability, and the 9 miscellaneous studies were assessed individually (Table 1).

**Table 1.**
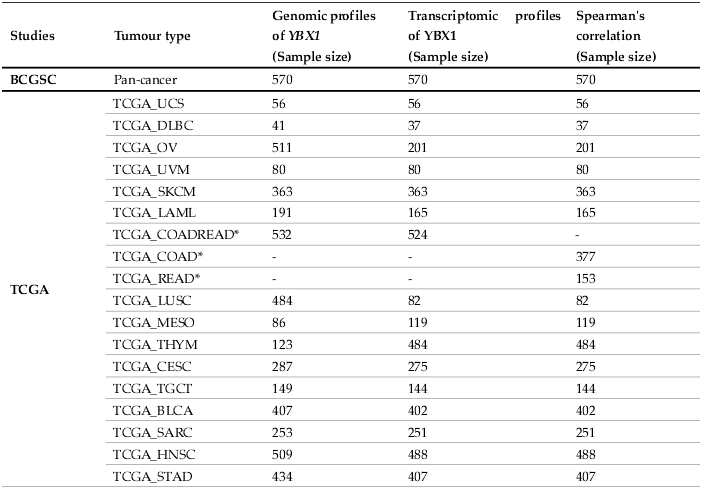
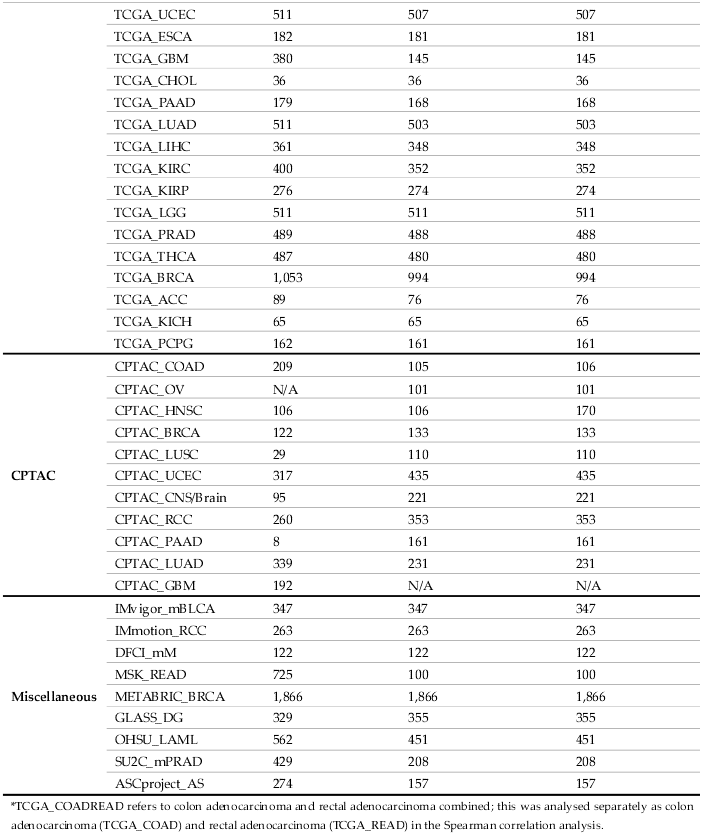
Summary of tumour types and sample sizes included in genomic and transcriptomic analyses of YBX1 and Spearman correlation analysis. N/A – not available.

The mRNA expression of *YBX1* analysed in this study were determined by RNA sequencing across all studies, except for the METABRIC_BRCA dataset, which was profiled using microarray. For *YBX1* mRNA expression, the BCGSC dataset was assessed at the pan-cancer level, the TCGA dataset was assessed across 32 individual cancer types, the CPTAC dataset was assessed across 10 out of 11 individual cancer types due to data availability, and the 9 miscellaneous studies were assessed individually (Table 1).

### 4.3 Spearman’s correlation analysis and Pathway analysis

The mRNA co-expression data against YBX1 mRNA was downloaded from cBioPortal by querying the YBX1 gene across the BCGSC pan-cancer dataset, the TCGA pan-cancer dataset, the CPTAC dataset, and across the 9 miscellaneous studies. The samples sizes for each dataset are detailed in Table 1.

To identify genes consistently correlated with *YBX1* expression across multiple datasets, we collected all available CSV files containing gene-level correlation data (Spearmans correlation coefficients and associated q-values) between YBX1 *mRNA* and other genes across independent cancer datasets. Data were imported and standardized using R (version 4.4.3) with the dplyr, tidyr, tibble, and pheatmap packages. Columns were harmonized across datasets to include Gene, Cytoband, Correlation (ρ), p-value (p), q-value (q), and Dataset identifiers. Only genes present in all datasets were retained for downstream analysis. For each gene, we determined the number of datasets in which the correlation with *YBX1* met the following criteria: positive correlation (ρ ≥ 0.3 and p ≤ 0.01) or negative correlation (ρ ≤ −0.3 and p ≤ 0.01). Genes meeting these thresholds in ≥80% of datasets were considered consistently positively or negatively correlated with *YBX1*, respectively. For visualization, correlation values for the filtered genes were arranged into a matrix with genes as rows and datasets as columns. Genes were ordered by mean correlation across datasets to highlight the most consistently associated genes. Side-bar annotations were added to indicate each gene’s mean correlation. Correlation values were visualized using colour gradients: negative correlations in blue, positive correlations from yellow to red, and values near zero in white. Heatmaps were generated with pheatmap, using hierarchical clustering for columns (datasets) and optional clustering for rows (genes), with row annotations for mean correlation values. To increase robustness of results, genes with missing values in any dataset were removed. The analysis workflow produced summary statistics, including the number of genes consistently positively or negatively correlated with *YBX1* across datasets.

Pathway enrichment analysis was subsequently conducted using the Reactome Pathways 2024 database through EnrichR [54], [55], [56], with pathways of a q-value ≤ 0.01 being considered significant.

### 4.4 Enrichment of driver mutations in high or low *YBX1* tumours and Pathway analysis

To assess the relationship between *YBX1* mRNA expression and gene-specific mutation rates across multiple tumour types, we analyzed mutation data from the 33 individual tumour types of the TCGA dataset. Tumours were stratified into high and low *YBX1* expression groups based on the 75th and 25th percentiles of *YBX1* mRNA expression within each tumour type, respectively. For each gene, the mutation rate was calculated in both groups, and a delta value (Δ = high − low) was computed to quantify enrichment of mutations in high-YBX1 tumours. Genes with missing mutation rates in either group were excluded from further analysis. The sample sizes for each cancer type are summarised in Table 2.

**Table 2.**
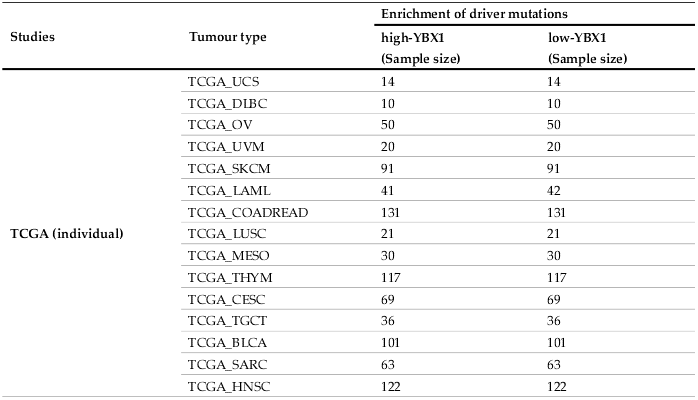
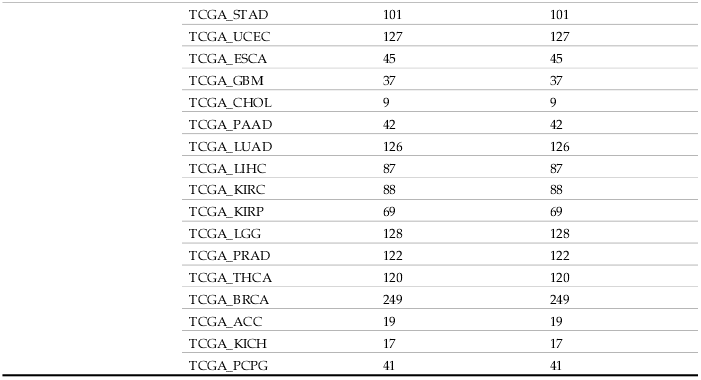
Summary of tumour types and sample sizes included in the enrichment of driver mutations in high or low YBX1 tumours.

| Studies | Tumour type | Enrichment of driver mutations |  |
| --- | --- | --- | --- |
|  |  | high-*YBX1*<br>(Sample size) | low-*YBX1*<br>(Sample size) |
| TCGA (individual) | TCGA_UCS | 14 | 14 |
|  | TCGA_DLBC | 10 | 10 |
|  | TCGA_OV | 50 | 50 |
|  | TCGA_UVM | 20 | 20 |
|  | TCGA_SKCM | 91 | 91 |
|  | TCGA_LAML | 41 | 42 |
|  | TCGA_COADREAD | 131 | 131 |
|  | TCGA_LUSC | 21 | 21 |
|  | TCGA_MESO | 30 | 30 |
|  | TCGA_THYM | 117 | 117 |
|  | TCGA_CESC | 69 | 69 |
|  | TCGA_TGCT | 36 | 36 |
|  | TCGA_BLCA | 101 | 101 |
|  | TCGA_SARC | 63 | 63 |
|  | TCGA_HNSC | 122 | 122 |

|  |  |  |
| --- | --- | --- |
| TCGA_STAD | 101 | 101 |
| TCGA_UCEC | 127 | 127 |
| TCGA_ESCA | 45 | 45 |
| TCGA_GBM | 37 | 37 |
| TCGA_CHOL | 9 | 9 |
| TCGA_PAAD | 42 | 42 |
| TCGA_LUAD | 126 | 126 |
| TCGA_LIHC | 87 | 87 |
| TCGA_KIRC | 88 | 88 |
| TCGA_KIRP | 69 | 69 |
| TCGA_LGG | 128 | 128 |
| TCGA_PRAD | 122 | 122 |
| TCGA_THCA | 120 | 120 |
| TCGA_BRCA | 249 | 249 |
| TCGA_ACC | 19 | 19 |
| TCGA_KICH | 17 | 17 |
| TCGA_PCPG | 41 | 41 |

A curated list of cancer-associated driver genes was obtained from previously published study by Bailey et al [57], and the analysis was restricted to these genes. Each gene was classified as Positive (Δ > 0), Negative (Δ < 0), or Unchanged (Δ = 0). For each gene, the number of tumour types with positive (pos) versus negative (neg) enrichment was summarized as a pos/neg ratio. Genes with a pos/neg ratio > 2 were considered consistently enriched in high-YBX1 tumours, whereas genes with a ratio ≤ 0.8 were considered depleted.

Mutation enrichment status across tumour types was visualized using heatmaps. Genes were arranged in rows and tumour types in columns, with Positive, Negative, Unchanged, or Unknown status encoded as discrete categories. To facilitate visualization, categorical status labels were mapped to numeric codes (Positive = 1, Negative = 2, Unchanged = 3, Unknown = 4), and heatmaps were generated using the pheatmap R package. Hierarchical clustering was applied to both genes and tumour types to reveal patterns of mutation enrichment across datasets. Separate heatmaps were produced for genes with pos/neg ratio > 2 and pos/neg ratio ≤ 0.8, highlighting consistent enrichment or depletion in high-YBX1 tumours.

All analyses were performed using R (version 4.4.3) with the tidyverse, pheatmap, dplyr, tidyr, readr, and tibble packages. Processed data and code for generating mutation delta values and heatmaps are available upon request. Gene ontology analysis was subsequently conducted using the GO Biological Processes 2025 database through EnrichR [54], [55], [56] with pathways of a q-value < 0.01 being considered significant. These are summarised in Supplementary Table S2.

### 4.5 Molecular characteristics associated with YBX1 mRNA levels

The molecular characteristics associated with *YBX1* expression levels were analysed using the BCGSC pan-cancer dataset, the TCGA pan-cancer dataset, and CPTAC studies across 10 out of 11 individual tumour types due to data availability (CPTAC_COAD, CPTAC_OV, CPTAC_HNSC, CPTAC_BRCA, CPTAC_LUSC, CPTAC_UCEC, CPTAC_CNS/Brain, CPTAC_RCC, CPTAC_PAAD, CPTAC_LUAD). The CPTAC studies were collectively assessed at the pan-cancer level and are hereafter referred to as the CPTAC pan-cancer dataset. Tumour samples in each pan-cancer dataset were stratified into high- and low-*YBX1* groups based on *YBX1* mRNA expression levels, with the high-*YBX1* group defined as the upper quartile (> 75th percentile) and the low-*YBX1* group defined as the lower quartile (< 25th percentile).

Molecular features analyzed included mutation count, fraction of genome altered (FGA), homologous recombination deficiency (HRD) score, and hypoxia scores (Ragnum, Winter, Buffa). Mutation count, capturing missense, insertion, deletion, and frameshift mutations, was available across all datasets. FGA, reflecting the proportion of the genome affected by copy number variations, was available only in TCGA, while HRD score, indicating homologous recombination repair defects, was available only in BCGSC. Together, these measures serve as surrogates for genomic stability. Comparisons between high- and low-*YBX1* tumours were performed using the Mann–Whitney U test, with p < 0.05 considered statistically significant.

Ragnum, Buffa, and Winter hypoxia scores are gene signature-based measure of transcriptional responses to low-oxygen levels within tumour microenvironments. This data was only available for the TCGA dataset. A Wilcoxon test was performed for comparisons between tumours with high and low YBX1, with a p-value < 0.05 considered to be statistically significant. The sample sizes for each molecular characteristics analysed are summarised in Table 3.

**Table 3.**
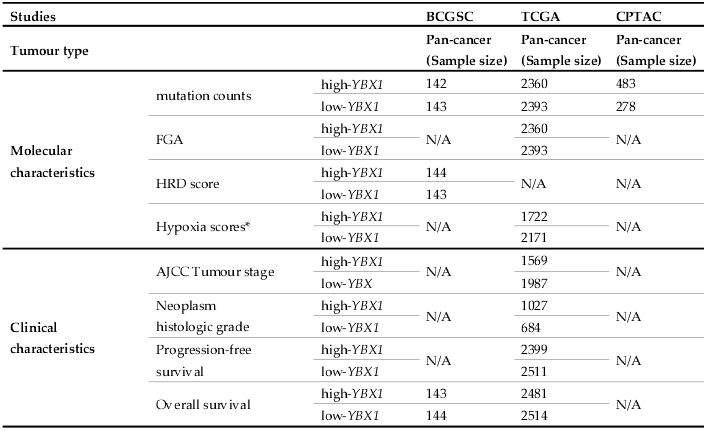
Summary of sample sizes included in the analyses of molecular and clinical characteristics associated with YBX1 mRNA levels.

| Studies |  |  | BCGSC | TCGA | CPTAC |
| --- | --- | --- | --- | --- | --- |
| Tumour type |  |  | Pan-cancer<br>(Sample size) | Pan-cancer<br>(Sample size) | Pan-cancer<br>(Sample size) |
| Molecular<br>characteristics | mutation counts | high-*YBX1* | 142 | 2360 | 483 |
|  |  | low-*YBX1* | 143 | 2393 | 278 |
|  | FGA | high-*YBX1* | N/A | 2360 | N/A |
|  |  | low-*YBX1* |  | 2393 |  |
|  | HRD score | high-*YBX1* | 144 | N/A | N/A |
|  |  | low-*YBX1* | 143 |  |  |
|  | Hypoxia scores* | high-*YBX1* | N/A | 1722 | N/A |
|  |  | low-*YBX1* |  | 2171 |  |
| Clinical<br>characteristics | AJCC Tumour stage | high-*YBX1* | N/A | 1569 | N/A |
|  |  | low-*YBX* |  | 1987 |  |
|  | Neoplasm<br>histologic grade | high-*YBX1* | N/A | 1027 | N/A |
|  |  | low-*YBX1* |  | 684 |  |
|  | Progression-free<br>survival | high-*YBX1* | N/A | 2399 | N/A |
|  |  | low-*YBX1* |  | 2511 |  |
|  | Overall survival | high-*YBX1* | 143 | 2481 | N/A |
|  |  | low-*YBX1* | 144 | 2514 |  |

### 4.6 Clinical characteristics associated with *YBX1* mRNA levels

Clinical characteristics associated with *YBX1* expression were analysed across the BCGSC and TCGA pan-cancer datasets by downloading the clinical data through cBioPortal. Tumour samples in each pan-cancer dataset were stratified into high- and low-*YBX1* groups based on *YBX1* mRNA expression levels, with the high-*YBX1* group defined as the upper quartile (> 75th percentile) and the low-*YBX1* group defined as the lower quartile (< 25th percentile).

The assessed parameters include the tumour stage derived from American Joint Committee on Cancer (AJCC) tumour staging system, neoplasm histologic grade, and patient survival status.

Tumour stage (T1-4) represents the size and extent of invasion of the primary tumour. Neoplasm histologic grade describes the degree of abnormality (low-, mid-, and high-grade) of the cancerous tissue. Due to data availability, tumour stage and grade were analysed using the TCGA dataset only. A Chi-squared test was performed for comparisons between tumours with high and low *YBX1*, with a p-value < 0.05 considered to be statistically significant. The sample sizes for each clinal characteristics analysed are summarised in Table 3.

Patient survival was assessed using both progression-free survival (available for the TCGA study only) and overall survival (available for the TCGA and BCGSC studies). The sample sizes for each clinal characteristics analysed are summarised in Table 3. Progression-free survival refers to the length of time during and following treatment that a patient survives without the disease worsening, whereas overall survival refers to the length of time from diagnosis or treatment initiation that patients survive, regardless of disease status. A log-rank test was performed for comparisons between tumours with high and low YBX1, with a p-value < 0.05 considered to be statistically significant.

## 5. Conclusions

In conclusion, our study defines a conserved high-*YBX1* tumour state characterized by genomic instability, transcriptional plasticity, and aggressive clinical behaviour. This pan-cancer framework unifies observations from single tumour-type studies and provides a foundation for mechanistic investigation and therapeutic targeting of high-YBX1 tumours across diverse cancer contexts.

## Supplementary Materials

**Figure S1:** Genomic alterations and YBX1 mRNA expression across additional independent datasets. **Table S1:** List of Reactome pathways enriched for mRNAs significantly correlated with YBX1 mRNA levels, based on EnrichR analysis. **Table S2:** Biological processes enriched among genes frequently mutated in high-YBX1 and low-YBX1 tumours, and those common to both, identified using EnrichR.

## Author Contributions

**SW, ZSP, and AB** performed experiments, analysed the data, **SW, and DS** assembled the data and contributed to the writing and reviewing the manuscript, **DS, GR, MS, AJ** and **AB’s** assisted with data interpretation and reviewed the manuscript, **SM** conceptualized and designed the study, interpreted, analysed and assembled the data, supervised the study, wrote and reviewed the manuscript.

## Institutional Review Board Statement

Not applicable.

## Informed Consent Statement

Not applicable.

## Data Availability Statement

Data presented in the study is available in publicly accessible repositories and the original data are openly available in cBioPortal [https://www.cbioportal.org/].

## Acknowledgments

None.

