## Supplementary figures and images for "Elevated YBX1 mRNA expression is associated with a genomically unstable and clinically aggressive cancer state: a pan-cancer analysis"

### Figure S1

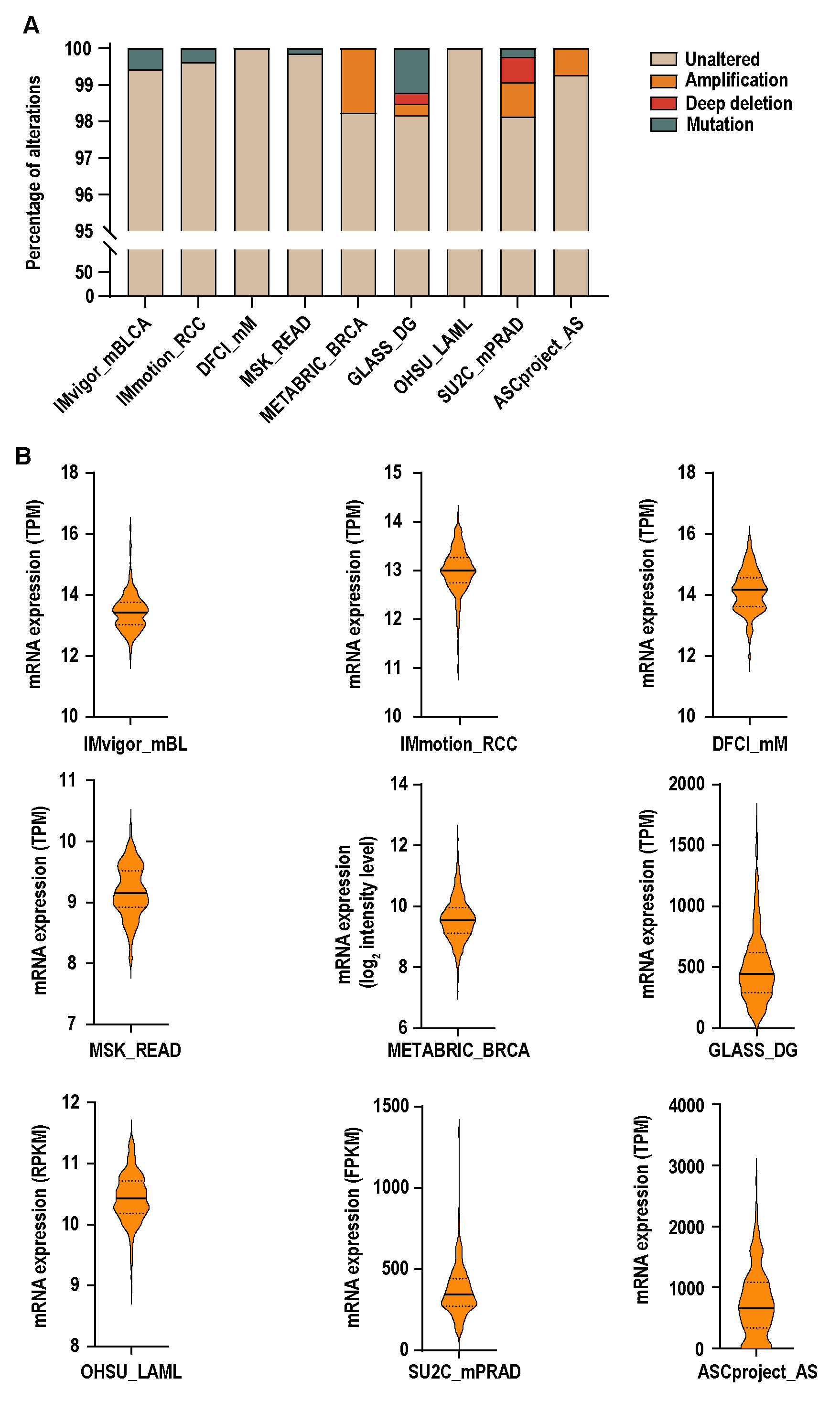
