## Supplementary material for "Elevated YBX1 mRNA expression is associated with a genomically unstable and clinically aggressive cancer state: a pan-cancer analysis": Table S1

Table S1. List of Reactome pathways enriched for mRNAs significantly correlated with *YBX1* mRNA levels based on EnrichR analysis.

| Term | P-value | Adjusted P-value |
| --- | --- | --- |
| Metabolism of RNA | 2.01E-10 | 5.17E-08 |
| mRNA Splicing - Major Pathway | 7.99E-05 | 1.28E-03 |
| mRNA Splicing | 9.79E-05 | 1.33E-03 |
| Processing of Capped Intron-Containing Pre-mRNA | 2.89E-04 | 3.39E-03 |
| Metabolism of Polyamines | 1.27E-03 | 6.33E-03 |
| AUF1 (hnRNP D0) Binds and Destabilizes mRNA | 1.06E-03 | 6.33E-03 |
| Major Pathway of rRNA Processing in the Nucleolus and Cytosol | 1.28E-03 | 6.33E-03 |
| rRNA Processing in the Nucleus and Cytosol | 1.48E-03 | 6.67E-03 |
| rRNA Processing | 2.44E-03 | 8.29E-03 |
| Mitotic Metaphase and Anaphase | 4.60E-06 | 3.96E-04 |
| Mitotic Anaphase | 4.50E-06 | 3.96E-04 |
| M Phase | 5.53E-05 | 1.10E-03 |
| Separation of Sister Chromatids | 4.60E-05 | 1.10E-03 |
| Regulation of APC C Activators Between G1 S and Early Anaphase | 6.34E-05 | 1.17E-03 |
| Regulation of Mitotic Cell Cycle | 8.50E-05 | 1.28E-03 |
| Cell Cycle, Mitotic | 2.57E-04 | 3.15E-03 |
| FBXL7 Down-Regulates AURKA During Mitotic Entry and in Early Mitosis | 1.06E-03 | 6.33E-03 |
| Asymmetric Localization of PCP Proteins | 1.56E-03 | 6.78E-03 |
| The Role of GTSE1 in G2 M Progression After G2 Checkpoint | 2.52E-03 | 8.33E-03 |
| SCF-beta-TrCP Mediated Degradation of Emi1 | 1.48E-05 | 9.55E-04 |
| Activation of APC C and APC C Cdc20 Mediated Degradation of Mitotic Proteins | 5.29E-05 | 1.10E-03 |
| APC C Cdc20 Mediated Degradation of Mitotic Proteins | 5.04E-05 | 1.10E-03 |
| APC C Cdh1 Mediated Degradation of Cdc20 and Other APC C Cdh1 Targets in Late Mitosis Early G1 | 4.58E-05 | 1.10E-03 |
| APC Cdc20 Mediated Degradation of Cell Cycle Proteins Prior to Satisfation of the Checkpoint | 4.58E-05 | 1.10E-03 |
| Cdc20 Phospho-APC C Mediated Degradation of Cyclin A | 4.36E-05 | 1.10E-03 |
| APC C Cdc20 Mediated Degradation of Securin | 3.35E-05 | 1.10E-03 |
| APC C-mediated Degradation of Cell Cycle Proteins | 8.50E-05 | 1.28E-03 |
| Ub-specific Processing Proteases | 1.30E-03 | 6.33E-03 |
| GLI3 Is Processed to GLI3R by the Proteasome | 1.33E-03 | 6.33E-03 |
| Degradation of GLI1 by the Proteasome | 1.33E-03 | 6.33E-03 |
| Degradation of GLI2 by the Proteasome | 1.33E-03 | 6.33E-03 |
| Degradation of DVL | 1.16E-03 | 6.33E-03 |
| GSK3B and BTRC CUL1-mediated-degradation of NFE2L2 | 1.06E-03 | 6.33E-03 |
| Degradation of AXIN | 1.06E-03 | 6.33E-03 |
| Vif-mediated Degradation of APOBEC3G | 1.06E-03 | 6.33E-03 |
| Vpu Mediated Degradation of CD4 | 9.61E-04 | 6.33E-03 |
| Autodegradation of the E3 Ubiquitin Ligase COP1 | 9.13E-04 | 6.33E-03 |
| Ubiquitin Mediated Degradation of Phosphorylated Cdc25A | 9.13E-04 | 6.33E-03 |
| Ubiquitin-dependent Degradation of Cyclin D | 9.13E-04 | 6.33E-03 |
| Regulation of Activated PAK-2p34 by Proteasome Mediated Degradation | 8.22E-04 | 6.33E-03 |
| Regulation of Ornithine Decarboxylase (ODC) | 8.67E-04 | 6.33E-03 |
| SCF(Skp2)-mediated Degradation of P27 P21 | 1.50E-03 | 6.67E-03 |
| Proteasome Assembly | 1.62E-03 | 6.78E-03 |
| Autodegradation of Cdh1 by Cdh1 APC C | 1.75E-03 | 6.82E-03 |
| Degradation of Beta-Catenin by the Destruction Complex | 3.08E-03 | 9.49E-03 |
| Regulation of Expression of SLITs and ROBOs | 3.42E-05 | 1.10E-03 |
| Signaling by ROBO Receptors | 8.94E-05 | 1.28E-03 |
| Axon Guidance | 3.11E-03 | 9.49E-03 |
| Cell Cycle Checkpoints | 2.06E-04 | 2.66E-03 |
| Stabilization of P53 | 1.16E-03 | 6.33E-03 |
| Regulation of Apoptosis | 9.61E-04 | 6.33E-03 |
| p53-Independent DNA Damage Response | 9.13E-04 | 6.33E-03 |
| p53-Independent G1 S DNA Damage Checkpoint | 9.13E-04 | 6.33E-03 |
| p53-Dependent G1 DNA Damage Response | 1.68E-03 | 6.78E-03 |
| p53-Dependent G1 S DNA Damage Checkpoint | 1.68E-03 | 6.78E-03 |
| G1 S DNA Damage Checkpoints | 1.81E-03 | 6.87E-03 |
| Host Interactions of HIV Factors | 3.94E-04 | 4.42E-03 |
| Cross-presentation of Soluble Exogenous Antigens (Endosomes) | 8.22E-04 | 6.33E-03 |
| Viral Infection Pathways | 8.67E-04 | 6.33E-03 |
| HIV Infection | 2.16E-03 | 7.53E-03 |
| Infectious Disease | 3.01E-03 | 9.49E-03 |
| Cell Cycle | 7.33E-04 | 6.33E-03 |
| Orc1 Removal From Chromatin | 2.01E-03 | 7.41E-03 |
| CDK-mediated Phosphorylation and Removal of Cdc6 | 2.15E-03 | 7.53E-03 |
| Cyclin E Associated Events During G1 S Transition | 3.16E-03 | 9.49E-03 |
| Cyclin A Cdk2-associated Events at S Phase Entry | 3.34E-03 | 9.89E-03 |
| NIK-->noncanonical NF-kB Signaling | 1.27E-03 | 6.33E-03 |
| Dectin-1 Mediated Noncanonical NF-kB Signaling | 1.44E-03 | 6.63E-03 |
| Activation of NF-kappaB in B Cells | 1.75E-03 | 6.82E-03 |
| Downstream Signaling Events of B Cell Receptor (BCR) | 2.75E-03 | 8.87E-03 |
| Hh Mutants Abrogate Ligand Secretion | 1.27E-03 | 6.33E-03 |
| Somitogenesis | 1.22E-03 | 6.33E-03 |
| Negative Regulation of NOTCH4 Signaling | 1.16E-03 | 6.33E-03 |
| Hh Mutants Are Degraded by ERAD | 1.11E-03 | 6.33E-03 |
| Hedgehog Ligand Biogenesis | 1.62E-03 | 6.78E-03 |
| Formation of Paraxial Mesoderm | 2.08E-03 | 7.45E-03 |
| Hedgehog 'On' State | 3.16E-03 | 9.49E-03 |
| Signaling by NOTCH4 | 3.08E-03 | 9.49E-03 |
| Oxygen-dependent Proline Hydroxylation of Hypoxia-inducible Factor Alpha | 1.68E-03 | 6.78E-03 |
| Cellular Response to Hypoxia | 2.29E-03 | 7.89E-03 |
| Regulation of RUNX3 Expression and Activity | 1.22E-03 | 6.33E-03 |
| Defective CFTR Causes Cystic Fibrosis | 1.38E-03 | 6.48E-03 |
| Regulation of RAS by GAPs | 1.81E-03 | 6.87E-03 |
| Regulation of PTEN Stability and Activity | 1.88E-03 | 7.01E-03 |
| Regulation of RUNX2 Expression and Activity | 2.08E-03 | 7.45E-03 |
| ABC Transporter Disorders | 2.52E-03 | 8.33E-03 |
| RNA Polymerase II Transcription Termination | 2.67E-03 | 8.73E-03 |
