## Supplementary material for "Elevated YBX1 mRNA expression is associated with a genomically unstable and clinically aggressive cancer state: a pan-cancer analysis": Table S2

Table S2. Biological processes enriched among genes frequently mutated in high-*YBX1* and low-*YBX1* tumours, and those common to both, identified using EnrichR.

| Term  (high *YBX1* exclusive) | P-value  (high *YBX1*) | Adjusted P-value (high *YBX1*) | P-value (low *YBX1*) | Adjusted P-value (low *YBX1*) |
| --- | --- | --- | --- | --- |
| Positive Regulation of Cell Migration (GO:0030335) | 3.52E-08 | 9.12E-06 | - | - |
| Chromatin Remodeling (GO:0006338) | 2.31E-07 | 4.26E-05 | - | - |
| Mammary Gland Epithelium Development (GO:0061180) | 4.60E-07 | 7.44E-05 | - | - |
| Positive Regulation of Cell Differentiation (GO:0045597) | 7.28E-07 | 1.05E-04 | - | - |
| Chromatin Organization (GO:0006325) | 1.56E-06 | 1.83E-04 | - | - |
| DNA Damage Response (GO:0006974) | 2.36E-06 | 2.54E-04 | - | - |
| Mismatch Repair (GO:0006298) | 3.48E-06 | 3.09E-04 | - | - |
| Cellular Response to Mechanical Stimulus (GO:0071260) | 3.57E-06 | 3.09E-04 | - | - |
| Negative Regulation of Transcription by RNA Polymerase II (GO:0000122) | 3.58E-06 | 3.09E-04 | - | - |
| Positive Regulation of RNA Biosynthetic Process (GO:1902680) | 5.58E-06 | 4.51E-04 | - | - |
| Negative Regulation of Programmed Cell Death (GO:0043069) | 6.90E-06 | 4.70E-04 | - | - |
| Regulation of Apoptotic Process (GO:0042981) | 1.20E-05 | 7.42E-04 | - | - |
| Enzyme-Linked Receptor Protein Signaling Pathway (GO:0007167) | 1.57E-05 | 9.21E-04 | - | - |
| Positive Regulation of Gene Expression (GO:0010628) | 2.17E-05 | 1.22E-03 | - | - |
| Positive Regulation of Macromolecule Biosynthetic Process (GO:0010557) | 2.63E-05 | 1.42E-03 | - | - |
| Regulation of Interleukin-1 Beta Production (GO:0032651) | 3.07E-05 | 1.53E-03 | - | - |
| Negative Regulation of Myoblast Differentiation (GO:0045662) | 4.42E-05 | 2.00E-03 | - | - |
| Regulation of Nitric Oxide Biosynthetic Process (GO:0045428) | 4.54E-05 | 2.00E-03 | - | - |
| Cellular Response to Oxygen-Containing Compound (GO:1901701) | 4.85E-05 | 2.00E-03 | - | - |
| Regulation of Fibroblast Proliferation (GO:0048145) | 4.96E-05 | 2.00E-03 | - | - |
| Development of Primary Male Sexual Characteristics (GO:0046546) | 4.96E-05 | 2.00E-03 | - | - |
| Male Gonad Development (GO:0008584) | 4.96E-05 | 2.00E-03 | - | - |
| Chromosome Organization (GO:0051276) | 5.40E-05 | 2.12E-03 | - | - |
| Regulation of Autophagy (GO:0010506) | 6.99E-05 | 2.58E-03 | - | - |
| Apoptotic Process (GO:0006915) | 7.38E-05 | 2.58E-03 | - | - |
| Vasculature Development (GO:0001944) | 7.58E-05 | 2.58E-03 | - | - |
| Maintenance of Gastrointestinal Epithelium (GO:0030277) | 7.58E-05 | 2.58E-03 | - | - |
| Mitotic Sister Chromatid Cohesion (GO:0007064) | 7.58E-05 | 2.58E-03 | - | - |
| Regulation of Cell Cycle (GO:0051726) | 7.98E-05 | 2.65E-03 | - | - |
| Positive Regulation of miRNA Metabolic Process (GO:2000630) | 8.69E-05 | 2.78E-03 | - | - |
| Positive Regulation of Cellular Process (GO:0048522) | 8.80E-05 | 2.78E-03 | - | - |
| Gonad Development (GO:0008406) | 9.35E-05 | 2.88E-03 | - | - |
| Regulation of Cytokine Production Involved in Inflammatory Response (GO:1900015) | 1.01E-04 | 2.99E-03 | - | - |
| Cellular Response to Nitrogen Compound (GO:1901699) | 1.02E-04 | 2.99E-03 | - | - |
| Positive Regulation of Interleukin-1 Beta Production (GO:0032731) | 1.08E-04 | 3.10E-03 | - | - |
| Negative Regulation of Cellular Process (GO:0048523) | 1.13E-04 | 3.14E-03 | - | - |
| Heart Morphogenesis (GO:0003007) | 1.16E-04 | 3.14E-03 | - | - |
| Positive Regulation of Interferon-Alpha Production (GO:0032727) | 1.19E-04 | 3.14E-03 | - | - |
| Mammary Gland Development (GO:0030879) | 1.19E-04 | 3.14E-03 | - | - |
| Regulation of Myoblast Differentiation (GO:0045661) | 1.24E-04 | 3.14E-03 | - | - |
| Peptidyl-Tyrosine Phosphorylation (GO:0018108) | 1.24E-04 | 3.14E-03 | - | - |
| Regulation of G1/S Transition of Mitotic Cell Cycle (GO:2000045) | 1.31E-04 | 3.25E-03 | - | - |
| Regulation of Interleukin-6 Production (GO:0032675) | 1.48E-04 | 3.57E-03 | - | - |
| Negative Regulation of Growth (GO:0045926) | 1.54E-04 | 3.57E-03 | - | - |
| Regulation of Lipid Storage (GO:0010883) | 1.56E-04 | 3.57E-03 | - | - |
| Positive Regulation of NLRP3 Inflammasome Complex Assembly (GO:1900227) | 1.56E-04 | 3.57E-03 | - | - |
| Positive Regulation of Catabolic Process (GO:0009896) | 1.57E-04 | 3.57E-03 | - | - |
| Positive Regulation of Interleukin-1 Production (GO:0032732) | 1.71E-04 | 3.78E-03 | - | - |
| Positive Regulation of Inflammasome-Mediated Signaling Pathway (GO:0141087) | 1.76E-04 | 3.78E-03 | - | - |
| G1/S Transition of Mitotic Cell Cycle (GO:0000082) | 1.82E-04 | 3.78E-03 | - | - |
| Regulation of Pentose-Phosphate Shunt (GO:0043456) | 1.90E-04 | 3.78E-03 | - | - |
| Reproductive System Development (GO:0061458) | 1.90E-04 | 3.78E-03 | - | - |
| Positive Regulation of Gonad Development (GO:1905941) | 1.90E-04 | 3.78E-03 | - | - |
| Positive Regulation of Male Gonad Development (GO:2000020) | 1.90E-04 | 3.78E-03 | - | - |
| Maintenance of DNA Repeat Elements (GO:0043570) | 1.90E-04 | 3.78E-03 | - | - |
| Negative Regulation of Cell Growth (GO:0030308) | 1.96E-04 | 3.84E-03 | - | - |
| Regulation of miRNA Transcription (GO:1902893) | 2.05E-04 | 3.87E-03 | - | - |
| Cell Cycle G1/S Phase Transition (GO:0044843) | 2.05E-04 | 3.87E-03 | - | - |
| Regulation of RNA Biosynthetic Process (GO:2001141) | 2.07E-04 | 3.87E-03 | - | - |
| Response to Cytokine (GO:0034097) | 2.11E-04 | 3.91E-03 | - | - |
| Regulation of Interferon-Alpha Production (GO:0032647) | 2.23E-04 | 4.06E-03 | - | - |
| Ras Protein Signal Transduction (GO:0007265) | 2.30E-04 | 4.07E-03 | - | - |
| Positive Regulation of Type I Interferon Production (GO:0032481) | 2.30E-04 | 4.07E-03 | - | - |
| Positive Regulation of Fibroblast Proliferation (GO:0048146) | 2.49E-04 | 4.35E-03 | - | - |
| Regulation of Translation (GO:0006417) | 2.61E-04 | 4.51E-03 | - | - |
| Regulation of Platelet Activation (GO:0010543) | 2.77E-04 | 4.59E-03 | - | - |
| Regulation of Tumor Necrosis Factor Production (GO:0032680) | 2.83E-04 | 4.59E-03 | - | - |
| Regulation of Male Gonad Development (GO:2000018) | 2.84E-04 | 4.59E-03 | - | - |
| Regulation of Osteoclast Development (GO:2001204) | 2.84E-04 | 4.59E-03 | - | - |
| Establishment of Mitotic Sister Chromatid Cohesion (GO:0034087) | 2.84E-04 | 4.59E-03 | - | - |
| Circulatory System Development (GO:0072359) | 2.93E-04 | 4.68E-03 | - | - |
| Gland Development (GO:0048732) | 3.35E-04 | 5.29E-03 | - | - |
| Regulation of Cytoskeleton Organization (GO:0051493) | 3.72E-04 | 5.75E-03 | - | - |
| DNA Repair (GO:0006281) | 3.73E-04 | 5.75E-03 | - | - |
| Mammary Gland Epithelial Cell Differentiation (GO:0060644) | 3.96E-04 | 6.03E-03 | - | - |
| Negative Regulation of Cell Differentiation (GO:0045596) | 4.23E-04 | 6.37E-03 | - | - |
| Regulation of Cell Growth (GO:0001558) | 4.44E-04 | 6.56E-03 | - | - |
| Regulation of Cell Cycle Phase Transition (GO:1901987) | 4.46E-04 | 6.56E-03 | - | - |
| Regulation of ERK1 and ERK2 Cascade (GO:0070372) | 4.55E-04 | 6.61E-03 | - | - |
| Positive Regulation of Tumor Necrosis Factor Production (GO:0032760) | 4.71E-04 | 6.77E-03 | - | - |
| Regulation of Interleukin-17 Production (GO:0032660) | 4.87E-04 | 6.84E-03 | - | - |
| Platelet Aggregation (GO:0070527) | 4.87E-04 | 6.84E-03 | - | - |
| Regulation of Interleukin-23 Production (GO:0032667) | 5.27E-04 | 7.03E-03 | - | - |
| Anoikis (GO:0043276) | 5.27E-04 | 7.03E-03 | - | - |
| Somatic Recombination of Immunoglobulin Gene Segments (GO:0016447) | 5.27E-04 | 7.03E-03 | - | - |
| Embryonic Digestive Tract Morphogenesis (GO:0048557) | 5.27E-04 | 7.03E-03 | - | - |
| Cellular Response to Growth Factor Stimulus (GO:0071363) | 5.27E-04 | 7.03E-03 | - | - |
| Positive Regulation of Tumor Necrosis Factor Superfamily Cytokine Production (GO:1903557) | 5.40E-04 | 7.13E-03 | - | - |
| Regulation of Endothelial Cell Migration (GO:0010594) | 6.16E-04 | 7.97E-03 | - | - |
| Regulation of Mitotic Cell Cycle Phase Transition (GO:1901990) | 6.16E-04 | 7.97E-03 | - | - |
| Embryonic Forelimb Morphogenesis (GO:0035115) | 6.75E-04 | 8.65E-03 | - | - |
| Regulation of Cytokine-Mediated Signaling Pathway (GO:0001959) | 7.23E-04 | 9.08E-03 | - | - |
| Regulation of DNA Recombination (GO:0000018) | 7.23E-04 | 9.08E-03 | - | - |
| Regulation of Inflammatory Response (GO:0050727) | 8.11E-04 | 9.96E-03 | - | - |
| Regulation of NLRP3 Inflammasome Complex Assembly (GO:1900225) | 8.35E-04 | 9.96E-03 | - | - |
| Regulation of Mesenchymal Stem Cell Differentiation (GO:2000739) | 8.42E-04 | 9.96E-03 | - | - |
| Cellular Response to Oxidised Low-Density Lipoprotein Particle Stimulus (GO:0140052) | 8.42E-04 | 9.96E-03 | - | - |
| Response to X-ray (GO:0010165) | 8.42E-04 | 9.96E-03 | - | - |
| Forelimb Morphogenesis (GO:0035136) | 8.42E-04 | 9.96E-03 | - | - |
| B Cell Activation (GO:0042113) | 8.54E-04 | 9.96E-03 | - | - |
| Regulation of Actin Filament-Based Process (GO:0032970) | 8.54E-04 | 9.96E-03 | - | - |
| Positive Regulation of Signal Transduction (GO:0009967) | 9.36E-04 | 1.08E-02 | - | - |
| Heart Development (GO:0007507) | 9.53E-04 | 1.08E-02 | - | - |
| Regulation of Microtubule-Based Process (GO:0032886) | 9.57E-04 | 1.08E-02 | - | - |
| Positive Regulation of Autophagy (GO:0010508) | 9.58E-04 | 1.08E-02 | - | - |
| Response to UV (GO:0009411) | 9.95E-04 | 1.11E-02 | - | - |
| Positive Regulation of Execution Phase of Apoptosis (GO:1900119) | 1.03E-03 | 1.12E-02 | - | - |
| Positive Regulation of Platelet Activation (GO:0010572) | 1.03E-03 | 1.12E-02 | - | - |
| Positive Regulation of Platelet Aggregation (GO:1901731) | 1.03E-03 | 1.12E-02 | - | - |
| Positive Regulation of MAPK Cascade (GO:0043410) | 1.09E-03 | 1.15E-02 | - | - |
| Peptidyl-Tyrosine Modification (GO:0018212) | 1.09E-03 | 1.15E-02 | - | - |
| Homotypic Cell-Cell Adhesion (GO:0034109) | 1.09E-03 | 1.15E-02 | - | - |
| Regulation of Defense Response (GO:0031347) | 1.11E-03 | 1.16E-02 | - | - |
| Negative Regulation of Gene Expression (GO:0010629) | 1.12E-03 | 1.16E-02 | - | - |
| Regulation of Actin Cytoskeleton Organization (GO:0032956) | 1.15E-03 | 1.16E-02 | - | - |
| Negative Regulation of Cold-Induced Thermogenesis (GO:0120163) | 1.16E-03 | 1.16E-02 | - | - |
| Stem Cell Differentiation (GO:0048863) | 1.16E-03 | 1.16E-02 | - | - |
| Positive Regulation of miRNA Transcription (GO:1902895) | 1.16E-03 | 1.16E-02 | - | - |
| Positive Regulation of Protein-Containing Complex Assembly (GO:0031334) | 1.19E-03 | 1.19E-02 | - | - |
| Axonal Fasciculation (GO:0007413) | 1.23E-03 | 1.20E-02 | - | - |
| Interleukin-6-Mediated Signaling Pathway (GO:0070102) | 1.23E-03 | 1.20E-02 | - | - |
| Regulation of Canonical NF-kappaB Signal Transduction (GO:0043122) | 1.23E-03 | 1.20E-02 | - | - |
| Protein Phosphorylation (GO:0006468) | 1.29E-03 | 1.25E-02 | - | - |
| Positive Regulation of Cell Population Proliferation (GO:0008284) | 1.31E-03 | 1.26E-02 | - | - |
| Positive Regulation of Translation (GO:0045727) | 1.32E-03 | 1.26E-02 | - | - |
| Small GTPase-mediated Signal Transduction (GO:0007264) | 1.45E-03 | 1.36E-02 | - | - |
| Positive Regulation of T Cell Activation (GO:0050870) | 1.46E-03 | 1.36E-02 | - | - |
| Regulation of Ubiquitin-Dependent Protein Catabolic Process (GO:2000058) | 1.47E-03 | 1.36E-02 | - | - |
| Epithelial to Mesenchymal Transition (GO:0001837) | 1.47E-03 | 1.36E-02 | - | - |
| Negative Regulation of Cytokine Production (GO:0001818) | 1.56E-03 | 1.41E-02 | - | - |
| Signal Transduction by P53 Class Mediator (GO:0072331) | 1.56E-03 | 1.41E-02 | - | - |
| Positive Regulation of Reactive Oxygen Species Metabolic Process (GO:2000379) | 1.56E-03 | 1.41E-02 | - | - |
| Mitotic Cell Cycle Phase Transition (GO:0044772) | 1.56E-03 | 1.41E-02 | - | - |
| B Cell Differentiation (GO:0030183) | 1.64E-03 | 1.47E-02 | - | - |
| Intrinsic Apoptotic Signaling Pathway (GO:0097193) | 1.67E-03 | 1.47E-02 | - | - |
| Embryonic Cranial Skeleton Morphogenesis (GO:0048701) | 1.68E-03 | 1.47E-02 | - | - |
| Regulation of RNA Export From Nucleus (GO:0046831) | 1.68E-03 | 1.47E-02 | - | - |
| Cellular Response to Cytokine Stimulus (GO:0071345) | 1.74E-03 | 1.51E-02 | - | - |
| Negative Regulation of Canonical Wnt Signaling Pathway (GO:0090090) | 1.78E-03 | 1.53E-02 | - | - |
| Regulation of Execution Phase of Apoptosis (GO:1900117) | 1.94E-03 | 1.62E-02 | - | - |
| Negative Regulation of Macrophage Activation (GO:0043031) | 1.94E-03 | 1.62E-02 | - | - |
| Positive Regulation of Cytoplasmic Translation (GO:2000767) | 1.94E-03 | 1.62E-02 | - | - |
| Genitalia Development (GO:0048806) | 1.94E-03 | 1.62E-02 | - | - |
| Mitotic Recombination (GO:0006312) | 1.94E-03 | 1.62E-02 | - | - |
| Cell Surface Receptor Protein Tyrosine Kinase Signaling Pathway (GO:0007169) | 1.96E-03 | 1.63E-02 | - | - |
| Negative Regulation of Macromolecule Biosynthetic Process (GO:0010558) | 1.99E-03 | 1.64E-02 | - | - |
| Cellular Response to Oxidative Stress (GO:0034599) | 2.01E-03 | 1.64E-02 | - | - |
| Cellular Response to Interleukin-1 (GO:0071347) | 2.03E-03 | 1.64E-02 | - | - |
| mRNA Stabilization (GO:0048255) | 2.03E-03 | 1.64E-02 | - | - |
| Eye Development (GO:0001654) | 2.13E-03 | 1.71E-02 | - | - |
| Response to UV-B (GO:0010224) | 2.21E-03 | 1.74E-02 | - | - |
| Digestive Tract Morphogenesis (GO:0048546) | 2.21E-03 | 1.74E-02 | - | - |
| Lipopolysaccharide-Mediated Signaling Pathway (GO:0031663) | 2.21E-03 | 1.74E-02 | - | - |
| Response to Interleukin-1 (GO:0070555) | 2.24E-03 | 1.76E-02 | - | - |
| Cellular Response to Lipopolysaccharide (GO:0071222) | 2.26E-03 | 1.76E-02 | - | - |
| Negative Regulation of MAPK Cascade (GO:0043409) | 2.40E-03 | 1.86E-02 | - | - |
| Negative Regulation of Canonical NF-kappaB Signal Transduction (GO:0043124) | 2.46E-03 | 1.90E-02 | - | - |
| Regulation of Canonical Wnt Signaling Pathway (GO:0060828) | 2.66E-03 | 2.04E-02 | - | - |
| Negative Regulation of Metabolic Process (GO:0009892) | 2.70E-03 | 2.06E-02 | - | - |
| Neuron Projection Guidance (GO:0097485) | 2.76E-03 | 2.07E-02 | - | - |
| Regulation of Response to External Stimulus (GO:0032101) | 2.76E-03 | 2.07E-02 | - | - |
| Positive Regulation of Cell Activation (GO:0050867) | 2.80E-03 | 2.07E-02 | - | - |
| Positive Regulation of Cytokine Production Involved in Inflammatory Response (GO:1900017) | 2.80E-03 | 2.07E-02 | - | - |
| Positive Regulation of Homotypic Cell-Cell Adhesion (GO:0034112) | 2.80E-03 | 2.07E-02 | - | - |
| Regulation of Phosphatidylinositol 3-Kinase/Protein Kinase B Signal Transduction (GO:0051896) | 2.82E-03 | 2.07E-02 | - | - |
| Positive Regulation of Mitotic Cell Cycle Phase Transition (GO:1901992) | 2.83E-03 | 2.07E-02 | - | - |
| Regulation of Endocytosis (GO:0030100) | 2.95E-03 | 2.11E-02 | - | - |
| Regulation of Extrinsic Apoptotic Signaling Pathway (GO:2001236) | 2.95E-03 | 2.11E-02 | - | - |
| Positive Regulation of T Cell Proliferation (GO:0042102) | 2.95E-03 | 2.11E-02 | - | - |
| Regulation of Cell-Matrix Adhesion (GO:0001952) | 2.95E-03 | 2.11E-02 | - | - |
| Embryonic Digestive Tract Development (GO:0048566) | 3.12E-03 | 2.17E-02 | - | - |
| Limb Morphogenesis (GO:0035108) | 3.12E-03 | 2.17E-02 | - | - |
| Mesoderm Development (GO:0007498) | 3.12E-03 | 2.17E-02 | - | - |
| Regulation of Behavior (GO:0050795) | 3.12E-03 | 2.17E-02 | - | - |
| Positive Regulation of Cytokine Production (GO:0001819) | 3.12E-03 | 2.17E-02 | - | - |
| Negative Regulation of Cytokine-Mediated Signaling Pathway (GO:0001960) | 3.22E-03 | 2.23E-02 | - | - |
| Regulation of Necroptotic Process (GO:0060544) | 3.45E-03 | 2.34E-02 | - | - |
| Regulation of Nuclear-Transcribed mRNA Catabolic Process, Deadenylation-Dependent Decay (GO:1900151) | 3.45E-03 | 2.34E-02 | - | - |
| Positive Regulation of G2/M Transition of Mitotic Cell Cycle (GO:0010971) | 3.45E-03 | 2.34E-02 | - | - |
| Regulation of Cell Cycle G2/M Phase Transition (GO:1902749) | 3.45E-03 | 2.34E-02 | - | - |
| Positive Regulation of Apoptotic Signaling Pathway (GO:2001235) | 3.64E-03 | 2.45E-02 | - | - |
| Cellular Response to Lipid (GO:0071396) | 3.74E-03 | 2.50E-02 | - | - |
| Phagocytosis (GO:0006909) | 3.79E-03 | 2.50E-02 | - | - |
| Response to Ionizing Radiation (GO:0010212) | 3.79E-03 | 2.50E-02 | - | - |
| Regulation of Transcription Regulatory Region DNA Binding (GO:2000677) | 3.81E-03 | 2.50E-02 | - | - |
| Regulation of Cell-Substrate Junction Assembly (GO:0090109) | 3.81E-03 | 2.50E-02 | - | - |
| Negative Regulation of Wnt Signaling Pathway (GO:0030178) | 3.95E-03 | 2.58E-02 | - | - |
| Regulation of Cold-Induced Thermogenesis (GO:0120161) | 4.15E-03 | 2.70E-02 | - | - |
| Fc-epsilon Receptor Signaling Pathway (GO:0038095) | 4.18E-03 | 2.70E-02 | - | - |
| Negative Regulation of Apoptotic Signaling Pathway (GO:2001234) | 4.26E-03 | 2.73E-02 | - | - |
| Negative Regulation of Multicellular Organismal Process (GO:0051241) | 4.39E-03 | 2.80E-02 | - | - |
| Negative Regulation of Osteoclast Differentiation (GO:0045671) | 4.56E-03 | 2.82E-02 | - | - |
| Cellular Response to Interleukin-6 (GO:0071354) | 4.56E-03 | 2.82E-02 | - | - |
| Positive Regulation of Cell Cycle G2/M Phase Transition (GO:1902751) | 4.56E-03 | 2.82E-02 | - | - |
| Eye Morphogenesis (GO:0048592) | 4.56E-03 | 2.82E-02 | - | - |
| Modulation by Host of Viral Genome Replication (GO:0044827) | 4.56E-03 | 2.82E-02 | - | - |
| Axon Guidance (GO:0007411) | 4.56E-03 | 2.82E-02 | - | - |
| Lymphocyte Differentiation (GO:0030098) | 4.59E-03 | 2.83E-02 | - | - |
| Positive Regulation of ERK1 and ERK2 Cascade (GO:0070374) | 4.67E-03 | 2.85E-02 | - | - |
| Regulation of Non-Canonical NF-kappaB Signal Transduction (GO:1901222) | 4.76E-03 | 2.85E-02 | - | - |
| Nucleosome Organization (GO:0034728) | 4.76E-03 | 2.85E-02 | - | - |
| Cellular Response to UV (GO:0034644) | 4.76E-03 | 2.85E-02 | - | - |
| Positive Regulation of Interleukin-6 Production (GO:0032755) | 4.76E-03 | 2.85E-02 | - | - |
| Positive Regulation of Lymphocyte Proliferation (GO:0050671) | 4.76E-03 | 2.85E-02 | - | - |
| Positive Regulation of Cell Motility (GO:2000147) | 4.87E-03 | 2.90E-02 | - | - |
| Cell Chemotaxis (GO:0060326) | 4.94E-03 | 2.92E-02 | - | - |
| Adaptive Imm Resp Based on Som Recomb of Imm Rcptrs Built Frm IgSF Domains (GO:0002460) | 4.96E-03 | 2.92E-02 | - | - |
| Metanephros Development (GO:0001656) | 4.96E-03 | 2.92E-02 | - | - |
| Regulation of Cell Cycle G1/S Phase Transition (GO:1902806) | 5.11E-03 | 2.99E-02 | - | - |
| Response to Tumor Necrosis Factor (GO:0034612) | 5.30E-03 | 3.08E-02 | - | - |
| Cytokine-Mediated Signaling Pathway (GO:0019221) | 5.38E-03 | 3.08E-02 | - | - |
| Negative Regulation of Non-Canonical NF-kappaB Signal Transduction (GO:1901223) | 5.38E-03 | 3.08E-02 | - | - |
| Cellular Response to Low-Density Lipoprotein Particle Stimulus (GO:0071404) | 5.38E-03 | 3.08E-02 | - | - |
| Positive Regulation of Interleukin-17 Production (GO:0032740) | 5.38E-03 | 3.08E-02 | - | - |
| Wound Healing (GO:0042060) | 5.48E-03 | 3.13E-02 | - | - |
| + Reg of Phosphatidylinositol 3-Kinase/Prot Kinase B Signal Transduction (GO:0051897) | 5.71E-03 | 3.20E-02 | - | - |
| Regulation of Cytoplasmic Translation (GO:2000765) | 5.81E-03 | 3.20E-02 | - | - |
| Negative Regulation of Protein Localization to Nucleus (GO:1900181) | 5.81E-03 | 3.20E-02 | - | - |
| Regulation of Platelet Aggregation (GO:0090330) | 5.81E-03 | 3.20E-02 | - | - |
| Blood Vessel Endothelial Cell Migration (GO:0043534) | 5.81E-03 | 3.20E-02 | - | - |
| Regulation of Actin Polymerization or Depolymerization (GO:0008064) | 5.81E-03 | 3.20E-02 | - | - |
| Regulation of Cell Size (GO:0008361) | 5.81E-03 | 3.20E-02 | - | - |
| Regulation of Interleukin-8 Production (GO:0032677) | 5.87E-03 | 3.20E-02 | - | - |
| Negative Regulation of Protein Phosphorylation (GO:0001933) | 5.87E-03 | 3.20E-02 | - | - |
| Regulation of T Cell Proliferation (GO:0042129) | 5.87E-03 | 3.20E-02 | - | - |
| Wound Healing, Spreading of Cells (GO:0044319) | 6.25E-03 | 3.40E-02 | - | - |
| Cellular Response to Tumor Necrosis Factor (GO:0071356) | 6.69E-03 | 3.55E-02 | - | - |
| Positive Regulation of Biosynthetic Process (GO:0009891) | 6.69E-03 | 3.55E-02 | - | - |
| Negative Regulation of Type II Interferon Production (GO:0032689) | 6.72E-03 | 3.55E-02 | - | - |
| Toll-Like Receptor 4 Signaling Pathway (GO:0034142) | 6.72E-03 | 3.55E-02 | - | - |
| Hematopoietic Progenitor Cell Differentiation (GO:0002244) | 6.72E-03 | 3.55E-02 | - | - |
| Brain Development (GO:0007420) | 6.75E-03 | 3.55E-02 | - | - |
| Regulation of Type II Interferon Production (GO:0032649) | 6.90E-03 | 3.60E-02 | - | - |
| Positive Regulation of Endothelial Cell Migration (GO:0010595) | 6.90E-03 | 3.60E-02 | - | - |
| DNA-templated Transcription Elongation (GO:0006354) | 7.19E-03 | 3.71E-02 | - | - |
| Transcription Elongation by RNA Polymerase II (GO:0006368) | 7.19E-03 | 3.71E-02 | - | - |
| mRNA Splice Site Recognition (GO:0006376) | 7.19E-03 | 3.71E-02 | - | - |
| Regulation of Epithelial to Mesenchymal Transition (GO:0010717) | 7.57E-03 | 3.83E-02 | - | - |
| Negative Regulation of Signal Transduction (GO:0009968) | 7.57E-03 | 3.83E-02 | - | - |
| Negative Regulation of Cytokine Production Involved in Inflammatory Response (GO:1900016) | 7.69E-03 | 3.83E-02 | - | - |
| Fc Receptor Signaling Pathway (GO:0038093) | 7.69E-03 | 3.83E-02 | - | - |
| Cell Surface Toll-Like Receptor Signaling Pathway (GO:0140895) | 7.69E-03 | 3.83E-02 | - | - |
| Positive Regulation of Calcium-Mediated Signaling (GO:0050850) | 7.69E-03 | 3.83E-02 | - | - |
| Positive Regulation of Catalytic Activity (GO:0043085) | 7.69E-03 | 3.83E-02 | - | - |
| Positive Regulation of Nitric Oxide Biosynthetic Process (GO:0045429) | 7.69E-03 | 3.83E-02 | - | - |
| Pyroptotic Inflammatory Response (GO:0070269) | 7.69E-03 | 3.83E-02 | - | - |
| Regulation of Osteoblast Differentiation (GO:0045667) | 7.80E-03 | 3.87E-02 | - | - |
| Regulation of Dendritic Spine Morphogenesis (GO:0061001) | 8.19E-03 | 3.99E-02 | - | - |
| Regulation of Macrophage Activation (GO:0043030) | 8.19E-03 | 3.99E-02 | - | - |
| Positive Regulation of ATP-dependent Activity (GO:0032781) | 8.19E-03 | 3.99E-02 | - | - |
| Positive Regulation of Nitric Oxide Metabolic Process (GO:1904407) | 8.19E-03 | 3.99E-02 | - | - |
| Positive Regulation of Immune Response (GO:0050778) | 8.51E-03 | 4.13E-02 | - | - |
| DNA Damage Response, Signal Transduction by P53 Class Mediator (GO:0030330) | 8.71E-03 | 4.19E-02 | - | - |
| Negative Regulation of Interleukin-6 Production (GO:0032715) | 8.71E-03 | 4.19E-02 | - | - |
| Central Nervous System Development (GO:0007417) | 8.87E-03 | 4.25E-02 | - | - |
| B Cell Receptor Signaling Pathway (GO:0050853) | 9.25E-03 | 4.42E-02 | - | - |
| Positive Regulation of Neuron Projection Development (GO:0010976) | 9.53E-03 | 4.53E-02 | - | - |
| Cellular Response to Chemical Stress (GO:0062197) | 9.79E-03 | 4.58E-02 | - | - |
| Regulation of Developmental Growth (GO:0048638) | 9.80E-03 | 4.58E-02 | - | - |
| Regulation of Protein Polymerization (GO:0032271) | 9.80E-03 | 4.58E-02 | - | - |
| Tumor Necrosis Factor-Mediated Signaling Pathway (GO:0033209) | 9.80E-03 | 4.58E-02 | - | - |
| Regulation of Ras Protein Signal Transduction (GO:0046578) | 9.80E-03 | 4.58E-02 | - | - |
| Negative Regulation of Intracellular Signal Transduction (GO:1902532) | 1.00E-02 | 4.67E-02 | - | - |
| Hemopoiesis (GO:0030097) | 1.01E-02 | 4.67E-02 | - | - |
| Regulation of p38MAPK Cascade (GO:1900744) | 1.04E-02 | 4.72E-02 | - | - |
| Response to Transforming Growth Factor Beta (GO:0071559) | 1.04E-02 | 4.72E-02 | - | - |
| Toll-Like Receptor Signaling Pathway (GO:0002224) | 1.04E-02 | 4.72E-02 | - | - |
| Macrophage Activation (GO:0042116) | 1.04E-02 | 4.72E-02 | - | - |
| Regulation of T Cell Mediated Cytotoxicity (GO:0001914) | 1.04E-02 | 4.72E-02 | - | - |
| Regulation of Intracellular Signal Transduction (GO:1902531) | 1.07E-02 | 4.88E-02 | - | - |
| Sister Chromatid Segregation (GO:0000819) | 1.09E-02 | 4.92E-02 | - |  |
| Embryonic Limb Morphogenesis (GO:0030326) | 1.09E-02 | 4.92E-02 | - | - |
| Positive Regulation of Epithelial Cell Migration (GO:0010634) | 1.09E-02 | 4.92E-02 | - | - |
| Term  (low *YBX1* exclusive) | **P-value**  **(high *YBX1*)** | **Adjusted P-value (high *YBX1*)** | **P-value (low *YBX1*)** | **Adjusted P-value (low *YBX1*)** |
| Intracellular Signaling Cassette (GO:0141124) | - | - | 7.99E-06 | 1.63E-03 |
| Phosphatidylinositol 3-Kinase/Protein Kinase B Signal Transduction (GO:0043491) | - | - | 2.17E-05 | 1.63E-03 |
| MAPK Cascade (GO:0000165) | - | - | 2.18E-05 | 1.63E-03 |
| Regulation of CD8-positive, Alpha-Beta T Cell Differentiation (GO:0043376) | - | - | 3.95E-05 | 2.19E-03 |
| Negative Regulation of Alpha-Beta T Cell Differentiation (GO:0046639) | - | - | 5.26E-05 | 2.44E-03 |
| Response to Peptide Hormone (GO:0043434) | - | - | 6.48E-05 | 2.73E-03 |
| Regulation of DNA-templated Transcription Initiation (GO:2000142) | - | - | 8.45E-05 | 3.01E-03 |
| Regulation of CD4-positive, Alpha-Beta T Cell Differentiation (GO:0043370) | - | - | 1.24E-04 | 4.09E-03 |
| Regulation of Peroxisome Proliferator Activated Receptor Signaling Pathway (GO:0035358) | - | - | 1.46E-04 | 4.15E-03 |
| Positive Regulation of Alpha-Beta T Cell Differentiation (GO:0046638) | - | - | 1.46E-04 | 4.15E-03 |
| Regulation of Fat Cell Differentiation (GO:0045598) | - | - | 1.55E-04 | 4.15E-03 |
| Positive Regulation of Protein Localization to Membrane (GO:1905477) | - | - | 1.61E-04 | 4.15E-03 |
| Response to Insulin (GO:0032868) | - | - | 2.02E-04 | 4.92E-03 |
| Regulation of Epithelial Cell Proliferation (GO:0050678) | - | - | 2.93E-04 | 6.78E-03 |
| Positive Regulation of Protein Targeting to Membrane (GO:0090314) | - | - | 3.54E-04 | 7.79E-03 |
| Regulation of Protein Targeting to Membrane (GO:0090313) | - | - | 5.56E-04 | 1.11E-02 |
| Regulation of Transcription by RNA Polymerase I (GO:0006356) | - | - | 8.57E-04 | 1.47E-02 |
| Negative Regulation of Fat Cell Differentiation (GO:0045599) | - | - | 9.13E-04 | 1.51E-02 |
| Positive Regulation of Establishment of Protein Localization (GO:1904951) | - | - | 1.03E-03 | 1.65E-02 |
| Semaphorin-Plexin Signaling Pathway (GO:0071526) | - | - | 1.36E-03 | 2.09E-02 |
| Positive Regulation of Myeloid Leukocyte Differentiation (GO:0002763) | - | - | 1.65E-03 | 2.45E-02 |
| Mitotic Spindle Assembly (GO:0090307) | - | - | 1.97E-03 | 2.68E-02 |
| Epidermal Growth Factor Receptor Signaling Pathway (GO:0007173) | - | - | 2.13E-03 | 2.82E-02 |
| Positive Regulation of Fat Cell Differentiation (GO:0045600) | - | - | 2.22E-03 | 2.86E-02 |
| Regulation of Cell Differentiation (GO:0045595) | - | - | 2.72E-03 | 3.41E-02 |
| Myeloid Cell Differentiation (GO:0030099) | - | - | 2.88E-03 | 3.50E-02 |
| ERBB Signaling Pathway (GO:0038127) | - | - | 2.98E-03 | 3.53E-02 |
| Epigenetic Regulation of Gene Expression (GO:0040029) | - | - | 3.61E-03 | 4.18E-02 |
| Positive Regulation of MAP Kinase Activity (GO:0043406) | - | - | 3.84E-03 | 4.33E-02 |
| Regulation of MAP Kinase Activity (GO:0043405) | - | - | 4.67E-03 | 4.81E-02 |
| Spindle Assembly (GO:0051225) | - | - | 4.67E-03 | 4.81E-02 |
| Positive Regulation of Protein Serine/Threonine Kinase Activity (GO:0071902) | - | - | 4.92E-03 | 4.85E-02 |
| Term  (high and low *YBX1* common) | **P-value**  **(high *YBX1*)** | **Adjusted P-value (high *YBX1*)** | **P-value (low *YBX1*)** | **Adjusted P-value (low *YBX1*)** |
| Positive Regulation of DNA-templated Transcription (GO:0045893) | 5.99E-14 | 7.76E-11 | 4.83E-06 | 1.63E-03 |
| Positive Regulation of Transcription by RNA Polymerase II (GO:0045944) | 3.48E-12 | 2.10E-09 | 4.27E-05 | 2.19E-03 |
| Regulation of Transcription by RNA Polymerase II (GO:0006357) | 4.87E-12 | 2.10E-09 | 2.15E-05 | 1.63E-03 |
| Regulation of Gene Expression (GO:0010468) | 2.30E-09 | 7.45E-07 | 1.45E-05 | 1.63E-03 |
| Negative Regulation of Apoptotic Process (GO:0043066) | 1.54E-07 | 3.32E-05 | 4.09E-03 | 4.41E-02 |
| Negative Regulation of DNA-templated Transcription (GO:0045892) | 1.54E-06 | 1.83E-04 | 3.72E-04 | 7.84E-03 |
| Regulation of Cell Population Proliferation (GO:0042127) | 6.30E-06 | 4.63E-04 | 5.78E-04 | 1.11E-02 |
| Negative Regulation of Cell Population Proliferation (GO:0008285) | 6.44E-06 | 4.63E-04 | 1.81E-03 | 2.54E-02 |
| Regulation of DNA-templated Transcription (GO:0006355) | 1.15E-05 | 7.42E-04 | 7.76E-05 | 2.99E-03 |
| Positive Regulation of Intracellular Signal Transduction (GO:1902533) | 2.78E-05 | 1.44E-03 | 2.47E-05 | 1.63E-03 |
| Negative Regulation of RNA Biosynthetic Process (GO:1902679) | 1.09E-03 | 1.15E-02 | 3.94E-03 | 4.35E-02 |
| Myeloid Leukocyte Differentiation (GO:0002573) | 4.26E-03 | 2.73E-02 | 4.80E-03 | 4.83E-02 |
| Cellular Response to Epidermal Growth Factor Stimulus (GO:0071364) | 6.72E-03 | 3.55E-02 | 6.99E-04 | 1.29E-02 |
| Regulation of Macromolecule Biosynthetic Process (GO:0010556) | 6.75E-03 | 3.55E-02 | 1.69E-03 | 2.45E-02 |
| Response to Epidermal Growth Factor (GO:0070849) | 8.19E-03 | 3.99E-02 | 8.57E-04 | 1.47E-02 |
